# Core Transcription Programs Controlling Injury-Induced Neurodegeneration of Retinal Ganglion Cells

**DOI:** 10.1101/2022.01.20.477004

**Authors:** Feng Tian, Yuyan Cheng, Songlin Zhou, Qianbin Wang, Aboozar Monavarfeshani, Kun Gao, Weiqian Jiang, Riki Kawaguchi, Qing Wang, Mingjun Tang, Ryan Donahue, Huyan Meng, Anne Jacobi, Jiani Yin, Xinyi Cai, Shane Hegarty, Joanna Stanicka, Phillip Dmitriev, Daniel Taub, Clifford J. Woolf, Joshua R. Sanes, Daniel H. Geschwind, Zhigang He

## Abstract

Neurodegenerative diseases are characterized by neuronal death and regenerative failure. However, gene regulatory programs governing how initial neuronal injuries lead to neuronal death remain poorly understood. In adult mice, optic nerve crush (ONC) injury, which severs all axons of retinal ganglion cells (RGCs), results in massive death of axotomized RGCs and regenerative failure of survivors. We performed an *in vivo* CRISPR/Cas9-based genome-wide screen of 1893 transcription factors (TFs) to seek repressors of RGC survival and axon regeneration following ONC. In parallel, we profiled the epigenetic and transcriptional landscapes of injured RGCs by ATAC-seq and RNA-seq to identify critical injury responsive TFs and their targets. Remarkably, these independent analyses converged on a set of four ATF/CEBP transcription factors – ATF3, ATF4, C/EBPγ and CHOP (Ddit3) – as critical regulators of survival. Further studies indicate that these TFs contribute to two pro-death transcriptional programs: ATF3/CHOP preferentially regulate pathways activated by cytokines and innate immunity, whereas ATF4/C/EBPγ regulate pathways engaged by intrinsic neuronal stressors. Manipulation of these TFs also protects RGCs in an experimental model of glaucoma, a prevalent disease in which RGCs die. Together, our results reveal core transcription programs that transform an initial axonal insult into a degenerative result and suggest novel strategies for treating neurodegenerative diseases.

## INTRODUCTION

Despite their vast clinical heterogeneity, most neurodegenerative diseases share certain pathological outcomes, the most devastating of which is neuronal death. Distal axon damage and dying back neuropathy have been observed in the early stage of many neurodegenerative diseases that lead to loss of neurons, including glaucoma, amyotrophic lateral sclerosis (ALS), and Alzheimer’s disease (Quigley, 1983, 2016; Calkins et al., 2017; Edwards, 2019). However, the pathways that lead from axonal insults to neuronal death remain incompletely understood (Perlson et al., 2010; Almasieh and Levin, 2017).

To identify the mechanisms by which damage to axons in the adult central nervous system (CNS) result in neurodegeneration, we took advantage of the optic nerve crush (ONC) model. In this model, 80% of retinal ganglion cells (RGCs) are lost two weeks after injury and there is virtually no spontaneous axon regeneration (Aguayo et al., 1991; Williams et al., 2020). Here, axonal injury is clearly the primary insult and extensive neurodegeneration, and regenerative failure are pathological outcomes. Previous candidate gene approaches have revealed several pathways that affect RGC survival and/or axon regeneration (Williams et al., 2020; Crair and Mason, 2016; Varadarajan et al., 2022), but a comprehensive view of the regulatory landscape governing this process is lacking.

Dissection of complex biological processes has often been advanced by unbiased large-scale genetic screening of many tissues including the CNS (Geschwind and Konopka, 2009; Parikshak et al., 2015; Kampmann et al., 2020). The clustered regularly interspaced short palindromic repeats (CRISPR) technology (Cong et al., 2013; Mali et al., 2013) provides a powerful means to conduct such screens *in vivo*. For analysis of ONC, intravitreal injection of AAV2 vectors efficiently transduces most RGCs with minimal effect on other cell types (Park et al., 2008). In addition, the massive and reproducible RGC loss and the short experimental duration of the ONC model allow for survival and regeneration to be analyzed efficiently. Since gene expression changes are key injury responses (Chandran, 2016, Moore et al., 2009, He and Jin, 2016; Hilton and Bradke, 2017, Winter et al., 2021; Varadarajan et al., 2022), we reasoned that an *in vivo* systematic screening of TFs would be a good starting point to identify the critical regulators of neuronal survival and/or axon regeneration.

An alternative approach to identifying TFs that control neuronal injury responses is epigenetic profiling. To control target gene expression, TFs need to bind to specific DNA sequence motifs, but the TF binding sites are not always in an accessible chromatin environment. Moreover, injury may significantly change the chromatin accessibility for TFs (Finelli et al., 2013; Gaub et al., 2010; Palmisano et al., 2019; Puttagunta et al., 2014). The assay of transposase-accessible chromatin using sequencing (ATAC-seq) is a high-throughput method that quantifies accessible chromatin genome-wide (Buenrostro et al., 2013; Buenrostro et al., 2015; Corces et al., 2017). Thus, integrative analysis of ATAC-seq and gene expression changes measured by RNA-sequencing (RNA-seq) in the same cell-type permits unbiased identification of injury-reactive TFs and their transcription targets.

In this study, we conducted multiple orthogonal, but complementary, functional genomic analyses, starting with a comprehensive, *in vivo* CRISPR screen of all known mammalian TFs. In parallel, we performed a RNA-seq coupled to an ATAC-seq analysis to define transcriptional networks and their drivers in intact and injured RGCs. Remarkably these different approaches converged on an overlapping set of TFs as critical regulators of neuronal injury responses, highlighting a core transcription program that mediates injury-induced neuronal degenerative outcomes. In a companion paper, Jacobi et al (accompany paper) utilized high-throughput single cell RNA-seq to analyze pathways downstream of several well-validated promoters of survival and regeneration (Park et al., 2008; Sun et al., 2011). Their results define molecular programs for survival, degeneration, and regeneration, providing complementary insights into these key aspects of neuronal injury responses.

## RESULTS

### Optimization of in vivo CRISPR screening

CRISPR has two major components: a guide RNA (sgRNA) and the CRISPR-associated endonuclease 9 (Cas9). By targeting a specific gene, the gRNA guides the non-specific Cas9 to desired DNA locations and introduces site-specific double strand breaks that generate insertion/deletion mutations through imprecise repair. Due to the high efficiency of generating sgRNAs, CRISPR has been utilized for genome-wide screens extensively in cultured cells (Shalem et al., 2014; 2015) and more recently in *in vivo* models (Jin et al., 2020; Wertz et al., 2020). To perform an *in vivo* loss-of-function screen for TFs that promote degeneration and/or inhibit axon regeneration, we utilized the RIKEN and TFCat databases (Kanamori et al., 2004; Fulton et al., 2009) to generate a non-redundant list of 1893 transcription regulators in the mouse genome. We further optimized each of the individual steps to establish an efficient functional screening platform (Figure 1A).

**Figure 1.**
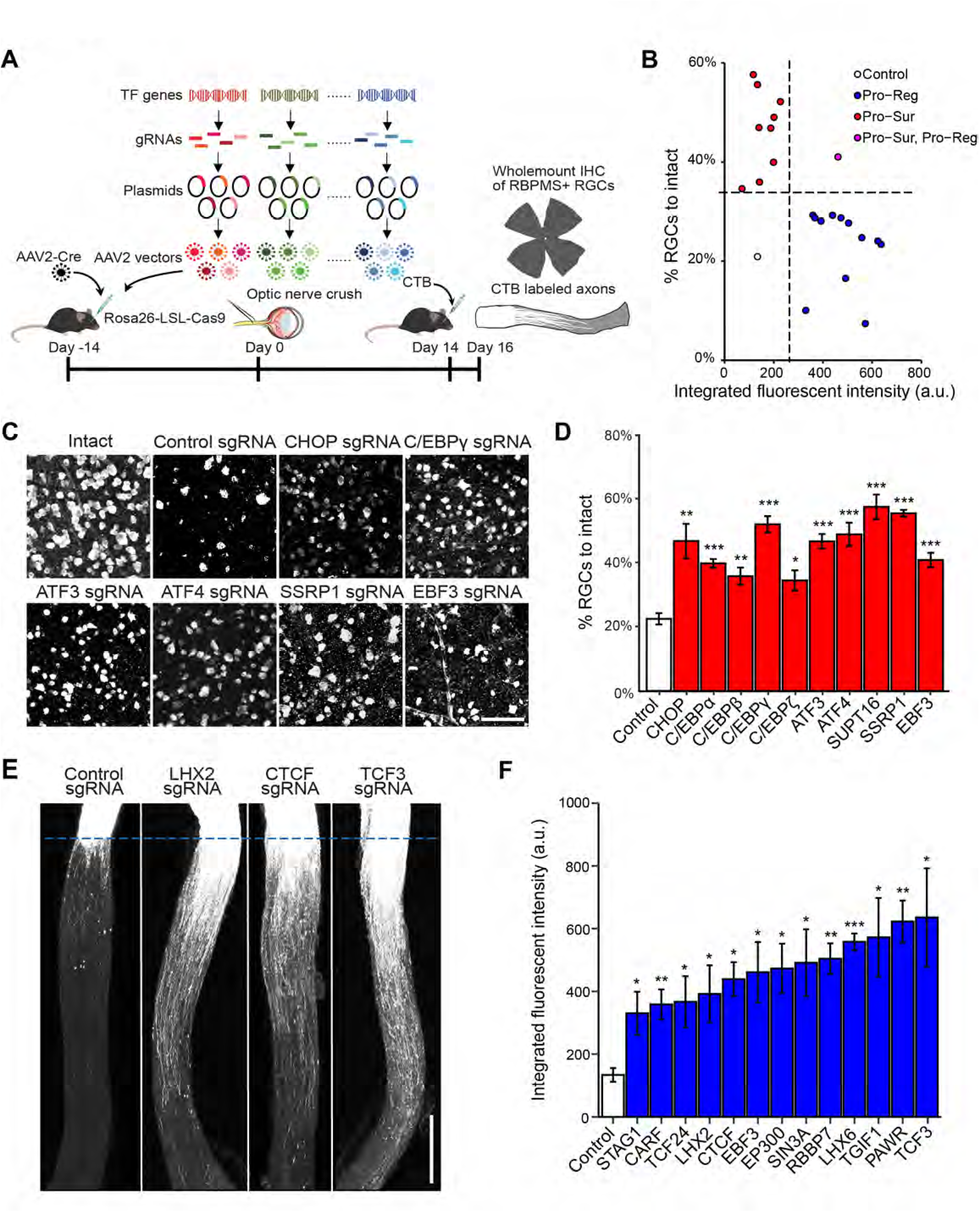
*In vivo* CRISPR screen identifies negative transcriptional regulators of RGC survival and axon regeneration after ONC. (A) Schematic illustration of the *in vivo* CRISPR screen in ONC model. AAV2 vectors encoding sgRNAs were injected intravitreally to Rosa26-LSL-Cas9 mice. To ensure ablation efficiency, the mix of five different sgRNAs (from GeCKO mouse v2 CRISPR knockout library) were selected to target each gene candidate. (B) The final lists of survival and regeneration hits were categorized into three groups: (1) deletion of the TF increasing axon regeneration without affecting RGC survival (Pro-Reg, 12 hits); (2) deletion of the TF increased RGC survival without affecting axon regeneration (Pro-Sur, 9 hits); (3) deletion of the TF promoted both RGC survival and axon regeneration (Pro-Sur, Pro-Reg, 1 hit). Each data point is the averaged results from RGC survival or axon regeneration analyses. (C) Representative immunohistochemistry images of wholemount retinas showing improved survival after ONC injury by CRISPR ablation of individual TFs. Scale bar, 50 μm. (D) Quantification of RGC survival with individual TF knockout. Data are shown as mean ± s.e.m. with n = 4-5 biological repeats. *p<0.05, **p<0.01, ***p<0.001. (E) Representative optic nerve images showing axon regeneration after ONC injury with individual TF knockout. Scale bar, 0.5 mm. (F) Quantification of CTB labeled fluorescent intensity (from crush site) for all sgRNA hits that promote RGC axon regeneration. Data are shown as mean ± s.e.m. with n = 4-6 biological repeats. *p<0.05, **p<0.01, ***p<0.001.

Briefly, to maximize CRISPR knockout efficiency, we selected five sgRNAs targeting different regions of each TF gene from the well-characterized genome-scale CRISPR/Cas9 knockout (GeCKO) library (Shalem et al., 2014; Shalem et al., 2015). Then, we generated a library of 1893 pools of AAV-sgRNA vectors, with each pool containing 5 different sgRNA-bearing AAV vectors targeting the same TF gene (Figure 1A). To introduce sgRNAs and Cas9 to RGCs, we co-injected pools of AAVs expressing sgRNAs and Cre into the vitreous bodies of Rosa26-LoxP-STOP-LoxP-Cas9-GFP (LSL-Cas9) mice. At 2 weeks after viral injection, ONC was performed, followed 2 weeks later by quantitative procedures to assess the extent of RGC survival and axon regeneration (Figure S1A).

As a proof-of-principle test, we showed that introducing sgRNAs for *Satb1*, an established marker for ON-OFF direction-selective RGCs (Peng et al., 2017), resulted in efficient Satb1 removal, as indicated by immunohistochemistry (Figure S1B and S1C). Similarly, with its specific sgRNAs, CRISPR mediated targeting of *Pten* gene also led to efficient reduction of PTEN expression (Figure S1D-E) and produced the expected increases in RGC survival and axon regeneration (Figure S1F-S1I) comparable to those seen in conventional PTEN knockout mice (Park et al., 2008).

### Identification of negative TF regulators of neuronal survival and/or axon regeneration

We then screened the effect of knockout of 1893 TFs, testing each gene in both eyes of one mouse. Genes whose knockout increased survival and/or regeneration (see Method) were retested in at least 3 additional eyes. Those producing consistent and significant increases in neuronal survival and/or axon regeneration were considered as true positive hits. This stringent screening protocol might miss TFs with modest effects (false negatives) but is expected to identify most, if not all, TFs with strong effects.

In total, from the 1893 TFs tested, we identified 10 genes as negative transcriptional regulators of neuronal survival (Figure 1B-D). Consistent with previous studies (Hu et al., 2012), CCCAAT/enhancer binding protein (C/EBP) homologous protein (CHOP, also called *Ddit3*), was identified as an anti-survival factor, providing additional verification of our screen method (Figure 1C and 1D). Other survival hits included several other C/EBP family members (C/EBPα, C/EBPβ, C/EBPγ, and CEBPζ) and two ATF family members (ATF3, ATF4), as well as two FACT complex components (SSRP1 and SUPT16), and EBF3, a tumor suppressor. We observed a similar increased survival of RGCs in ATF3 knockout mice (Renthal et al., 2020), validating the phenotype observed with the CRISPR experiment (Figure S1J-M). Notably, the identified ATFs and C/EBPs are in the basic leucine zipper domain (bZIP) containing proteins family. While these evolutionarily conserved TFs have been implicated in regulating several biological processes, such as memory inhibition (Bartsch et al., 1995; Abel et al., 1998, Sidrauski et al. 2013) and the integrated stress response (Wortel et al., 2017; Costa-Mattioli and Walter, 2020), the predominance of ATF and C/EBP members in our survival hits were a surprise, as they were not previously identified in this capacity in the literature.

Although our focus here is on neuronal survival, we also assessed axon regeneration and identified 13 genes as negative regulators of this process (Figure 1B, E, F). This group included several genes implicated in epigenetic regulation (CTCF/STAG1, histone acetyltransferase EP300, RBBP7, and SIN3A), members of the BHLH family (TCF3, TCF24) and the homeobox family (LHX2, LHX6, TGIF1) genes, tumor suppressors PAWR/WT1, and EBF3 (Figure 1E and 1F). The only overlapping gene among the survival and regeneration lists is EBF3. This dissociation suggests that transcription programs for regulating neuronal survival and axon regeneration are largely separate; the companion paper based primarily on transcriptomics (Jacobi et al., accompanying paper) describes these distinct programs in detail.

### Characterization of chromatin accessibility changes in RGCs following optic nerve crush

As a drastic insult, axotomy may trigger large-scale changes in gene expression via reorganization of chromatin. To assess genome-wide changes in chromatin accessibility following ONC, we performed ATAC-seq, which enables high-throughput quantification of open chromatin regions, as well as TF footprinting (Buenrostro et al., 2013; Bentson et al., 2020). In tandem, we also performed bulk RNA sequencing, allowing us to directly assess whether identified chromatin alterations led to transcriptional changes of the TFs and their target genes. We optimized purification of the RGCs from intact or injured mice for the ATAC-seq and RNA-seq studies (Figure 2A). Based on published data (Tran et al., 2019), we reasoned that most critical transcriptional responses to injury would occur in the first few days after injury, when gene expression has already been altered, but RGC death has not started. Thus, we FACS-sorted RGCs at 0, 1 or 3 days after injury and subjected them to ATAC-seq and RNA-seq (Figure 2A).

**Figure 2.**
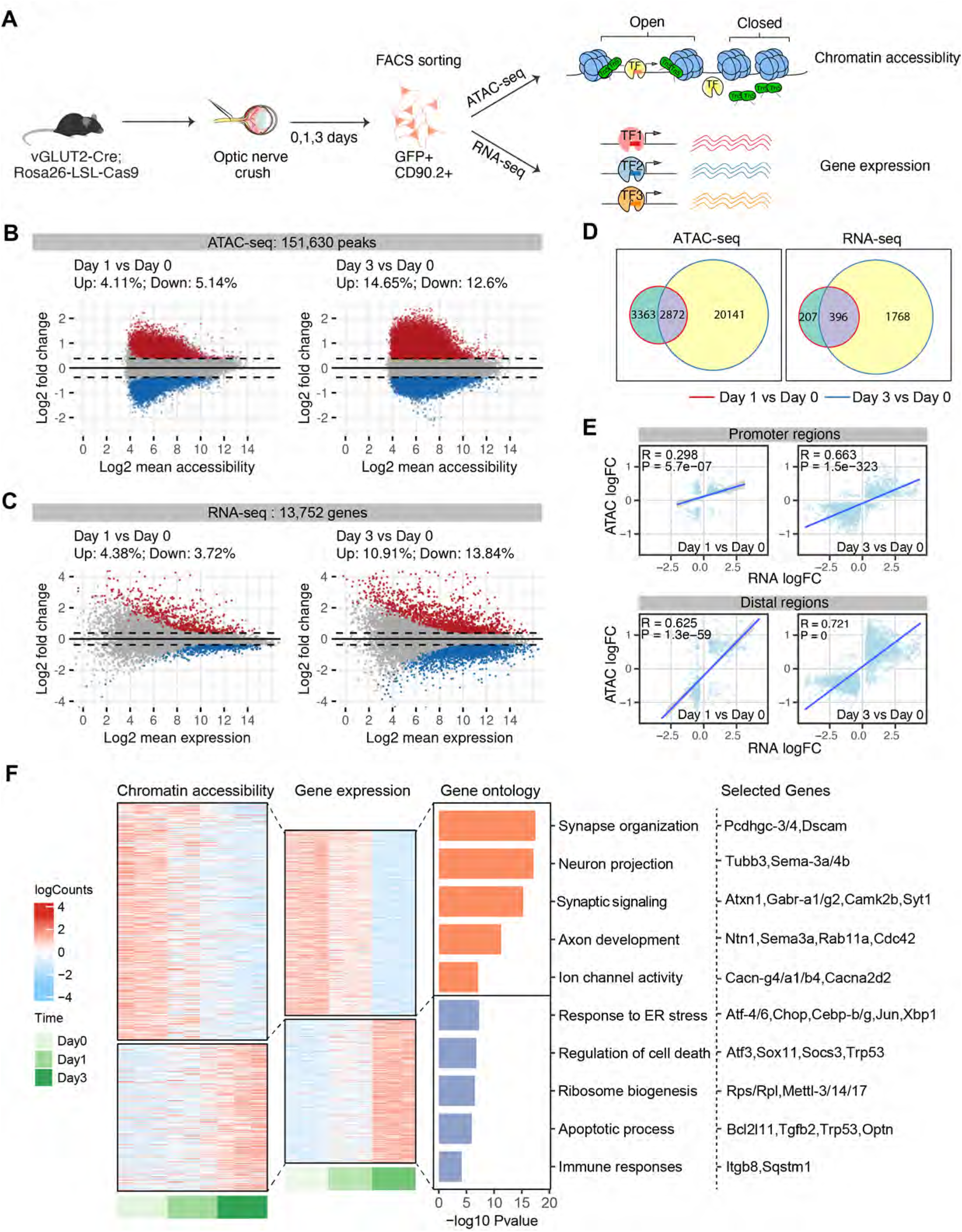
Characterization of chromatin accessibility changes in retinal ganglion cells following optic nerve crush. (A) Schematic diagram summarizing the overall experimental flow. vGLUT2-labeled RGCs were FACS-sorted at 0 (no crush) 1, 3 days following optic nerve crush. ATAC-seq and RNA-seq were performed on separate sets of injured RGCs. (B-D) MA plots displaying differential accessible regions (DAR) (B) and differential expressed genes (DEGs) (C) in RGCs following injury. Each dot represents a peak region or a gene, and colored dots indicate DARs or DEGs (FDR p < 0.1, | log2 FC| > 0.3). Up- regulated: red; Down-regulated: blue. (D) Overlap of DARs and DEGs at day 1 or day 3 following injury. (E) Pearson correlations between injury-induced changes in gene expression and chromatin accessibility at the promoter and distal DARs. Using GENCODE annotations, we defined an ATAC-seq peak ± 2 kb of a gene’s transcription start site (TSS) as a promoter (proximal regulatory element), and non-promoter peaks ± 500 kb of TSS as distal regulatory regions of that gene. The differential accessibilities of these DARs (log2 FCs) were correlated and plotted against the differential expression of the linked genes. If several distal peaks are linked to the same gene, the average differential accessibility was used to correlate with differential expression. (F) Chromatin accessibility and mRNA expression of linked peak-gene pairs. Heatmap colors indicate row-scaled chromatin accessibility (left) or mRNA expression levels (right). Bar plots represent negative log10 FDR-corrected p-values of top Gene Ontology (GO) terms associated with genes in each cluster. Representative genes in each term were displayed.

Following extensive quality control procedures (Figure S2A-G), we identified 151,630 reproducible accessible regions, or peaks, across the pooled ATAC-seq dataset. Importantly, we observed high correlations between biological replicates within peaks (Pearson’s correlation coefficient *r >* 0.95, Figure S2E). By analyzing chromatin accessibility changes in response to ONC, we identified 14,024 (∼9.25%) differentially accessible regions (DARs) at day 1 post-injury and 41,346 (∼27.25%) DARs at day 3 (Figure 2B, Table S1), with similar numbers of peaks bearing increased or reduced accessibility at both time points. Consistent with the ATAC-seq data, RNA-seq analysis on injured RGCs identified more changes at day 3 versus day 1: ∼ 8.1% (1,115) differentially expressed genes (DEGs) on day 1, and ∼ 24.75% DEGs (3,403) at day 3 (Figure 2C, Table S2). Only ∼50% of DARs or DEGs at day 1 overlap with those identified at day 3 (Figure 2D). Thus, these results revealed injury-induced, time-dependent alterations in both genomic accessibility and gene expression patterns in RGCs.

DARs are enriched in more accessible chromatin regions that were previously annotated from mouse brain (Ernst and Kellis, 2012; Yin et al., 2012), including, but not confined to promoters and enhancers and depleted in heterochromatin and transcribed regions that are less accessible (Figure S2C), in agreement with previous findings from other regions and cells (De La Torre Ubieta et al. 2018; Ernst and Kellis, 2012; Klemm et al., 2019; Shen et al., 2012). As the accessibility of both proximal promoter and distal regions such as enhancers is important for regulating gene expression (Roadmap Epigenomics et al., 2015; Spitz and Furlong, 2012), we further analyzed the relationships between gene expression and local chromatin accessibility patterns in our dataset (Figure 2E). The correlations across DEGs and DARs (both proximal and distal), increase at day 3 after injury compared to day 1, consistent with the expectation that changes in chromatin accessibility precede changes in gene expression. We also found that accessibility of distal DARs correlates more strongly with gene expression than promoter-proximal chromatin accessibility, a finding also observed in other cell types (Trevino et al., 2019).

We next identified significantly correlated peak-gene pairs (Table S3), defining two patterns of chromatin accessibility and gene expression in response to CNS injury (Figure 2F). The first cluster of peak-gene pairs includes those whose chromatin accessibility and gene expression are both down-regulated by injury. Functionally they are associated with synapse organization, synaptic transmission, cell adhesion and neuron projection (Figure 2F, Table S3). Among them are *Pcdhgc3/4* and *Dscam,* which have been implicated in RGC survival (Chen et al., 2012; Keeley et al., 2012). The second is an injury up-regulated cluster, which is involved in the regulation of endoplasmic reticulum stress, ribosome biogenesis, cell death / apoptosis, immune processes, and RNA processing (Figure 2F, Table S3). Interestingly, in addition to several genes previously implicated in stress/injury response and neuronal death, such as Sox11, Socs3, Gadd45g (Fischer et al., 2004; Sun et al. 2011, Norsworthy et al., 2017) and glaucoma-associated gene Optn (Wiggs and Pasquale, 2017), this cluster also includes several death-promoting TF hits identified from our CRISPR screening, including *Atf3, Atf4, CHOP, Cebpb, Cebpγ* (Figure 1, Figure 2F). Thus, these findings suggest that CNS injury induces widespread chromatin accessibility changes driving the expression of genes that are less favorable for neuronal survival and axon regrowth.

### Identification of TFs driving chromatin accessibility changes

To identify TFs whose expression was significantly correlated with changes in the accessibility of their binding motifs, we integrated ATAC-seq, RNA-seq and databases of TF-binding specificity. This approach is based on the rationale that the abundance of a TF and the accessibility of its binding motifs are associated with its gene regulatory activity (Corces et al., 2020; Trevino et al., 2020) (Figure 3A). By scanning DARs in the ATAC-seq data for TF motif occurrence, we identified a total of 1,310 significantly enriched TF binding motifs (Table S4). We than used ChromVAR (Schep et al., 2017) to compute changes in motif accessibility across injury conditions as deviation Z-scores, as an indicator of a TF’s activity at that position, which was further correlated with each TF’s mRNA expression from RNA-seq data (Table S4). Remarkably, among the top quartile of all TFs whose binding motifs showed the largest correlation between changes in their expression levels and accessibility of their binding motifs were four of the CRISPR screen hits ATF3, ATF4, C/EBPγ, CHOP/Ddit3 (Figure 3B, absolute Pearson correlations *r* > 0.5 and *P* < 0.05). mRNA and protein expression levels of all 4 TFs were consistently increased at day 3, as was the increased accessibility of their binding sites (Figure 3C and D).

**Figure 3.**
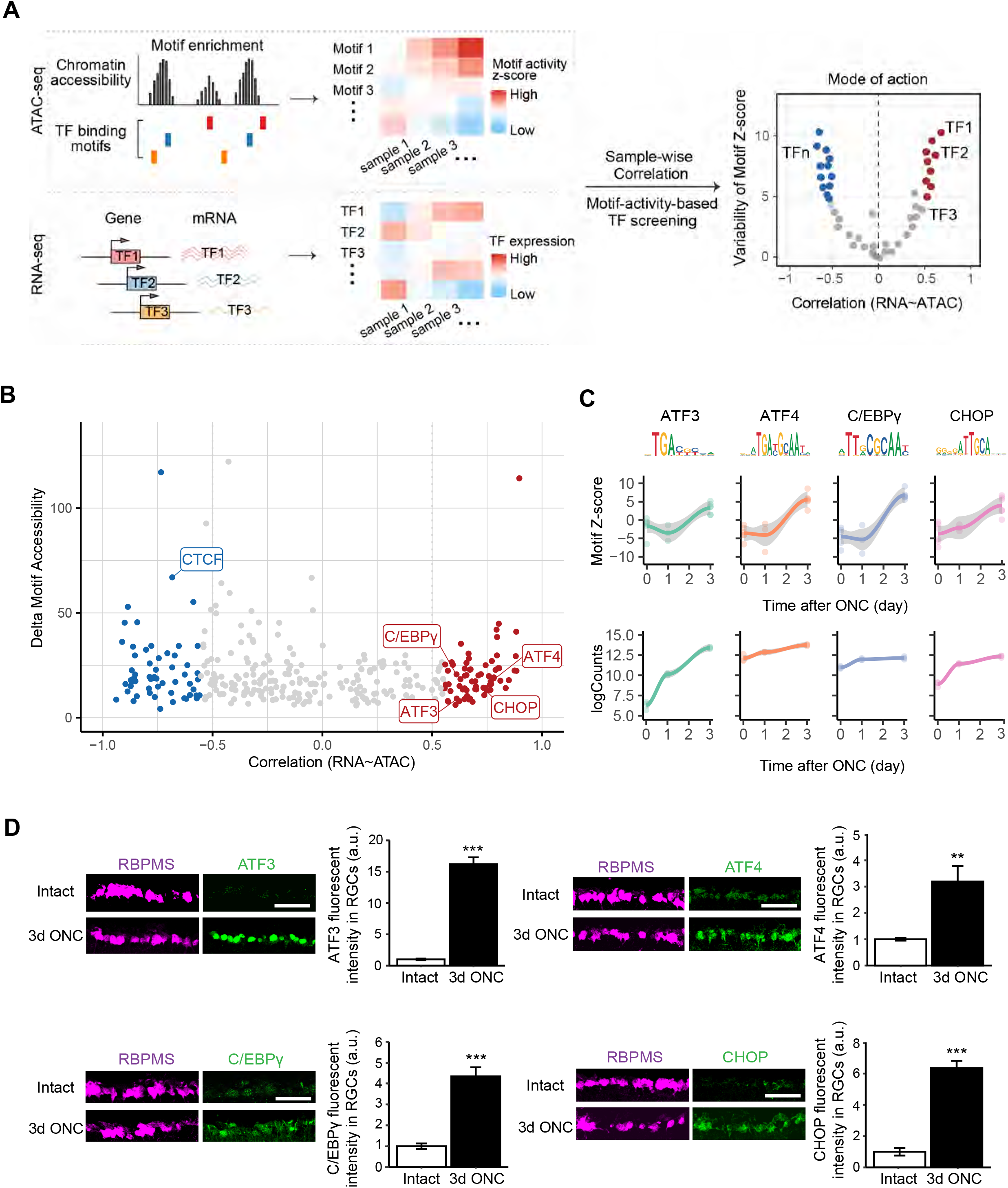
Identification of transcription factors driving chromatin accessibility changes. (A) A schematic diagram displaying an unbiased bioinformatic approach to identify TF regulators driving chromatin accessibility and gene expression changes in injured RGCs. In this approach, we first find TF binding motifs that are significantly enriched within differentially accessible regions (DAR). The degree of accessibility at enriched TF motifs was computed as deviation Z-scores, and were correlated with TF expression levels across samples to classify TF mode of actions. (B) TF gene expression - motif accessibility correlations against maximum inter-sample difference in deviation Z-score. Each dot represents a TF, and colored dots indicate TFs whose gene expressions are significantly correlated or anti-correlated to motif deviations (Pearson’s correlation coefficient *r* > 0.5 and *FDR* < 0.1), and whose maximum cross- sample difference in deviation z-score is in the top 25% of all TFs. Th genes overlapped with CRISPR screen hits were highlighted. (C) TF binding motif accessibility deviations and RNA-seq expression levels for ATF3, ATF4, C/EBPγ, CHOP/DDIT3. (D) Representative immunohistochemical images showing injury-induced protein expression at 3d post-ONC of individual TFs. Data are shown as mean ± s.e.m. with n = 4. **p<0.01, ***p<0.001. Scale bar, 10 μm.

### Characterization of functional TF targets by combinatorial analysis of DNA footprinting and RNA-seq of perturbed RGCs

For further computational and functional analysis we focused on a set of four genes – ATF3, ATF4, C/EBPγ and CHOP/Ddit3 – for several reasons. First, all four were hits in our CRISPR screen (Figure 1D). Second, these genes exhibited both increased chromatin accessibility and gene expression (Figures 2F and 3C). Third, ATAC-Seq data analysis implicated these genes as critical TFs that drive injury-induced responses in RGCs. (Figure 3B). In contrast, other hits from the CRISPR screen (SUPT16, SSRP1) are expressed in intact and injured RGCs with similar levels (Tran et al., 2019), suggesting a permissive role in injury responses. Finally, these genes, which are members of an evolutionarily conserved family, may act synergistically via dimerization to regulate gene expression programs (Reinke et al., 2013). Together, these data support the hypothesis that this family of TFs act as critical transcriptional regulators of early injury responses in RGCs.

To characterize survival regulating pathways, we used two complementary methods to identify the downstream targets of ATF3/4, C/EBPγ and CHOP in injured RGCs (Figure 4A). First, we characterized TF binding to target genes via DNA footprinting. Since TF binding shields bound DNA elements from transposase-mediated digestion, protected DNA sequences are believed to be direct TF binding sites, or footprints (Figure S3). Using newly developed methods (Bentson et al., 2020, Vierstra et al., 2020; Funk et al., 2020), which identify direct TF-DNA binding events (similar to ChIP-seq), we mapped genome-wide DNA footprints overlapping with each TF’s binding motifs, and quantified TF binding activities by leveraging ATAC-seq measurements of footprinted regions (Figure S3A). Notably, all four TFs become more active following injury, with increased binding depth and more open surrounding chromatin (Figure S3B-D), supporting their roles in driving injury-induced chromatin state changes. Mapping each TF’s footprints to individual genes revealed a highly significant correlation between footprint activities and gene expression changes induced by injury (Figure S3E),

**Figure 4.**
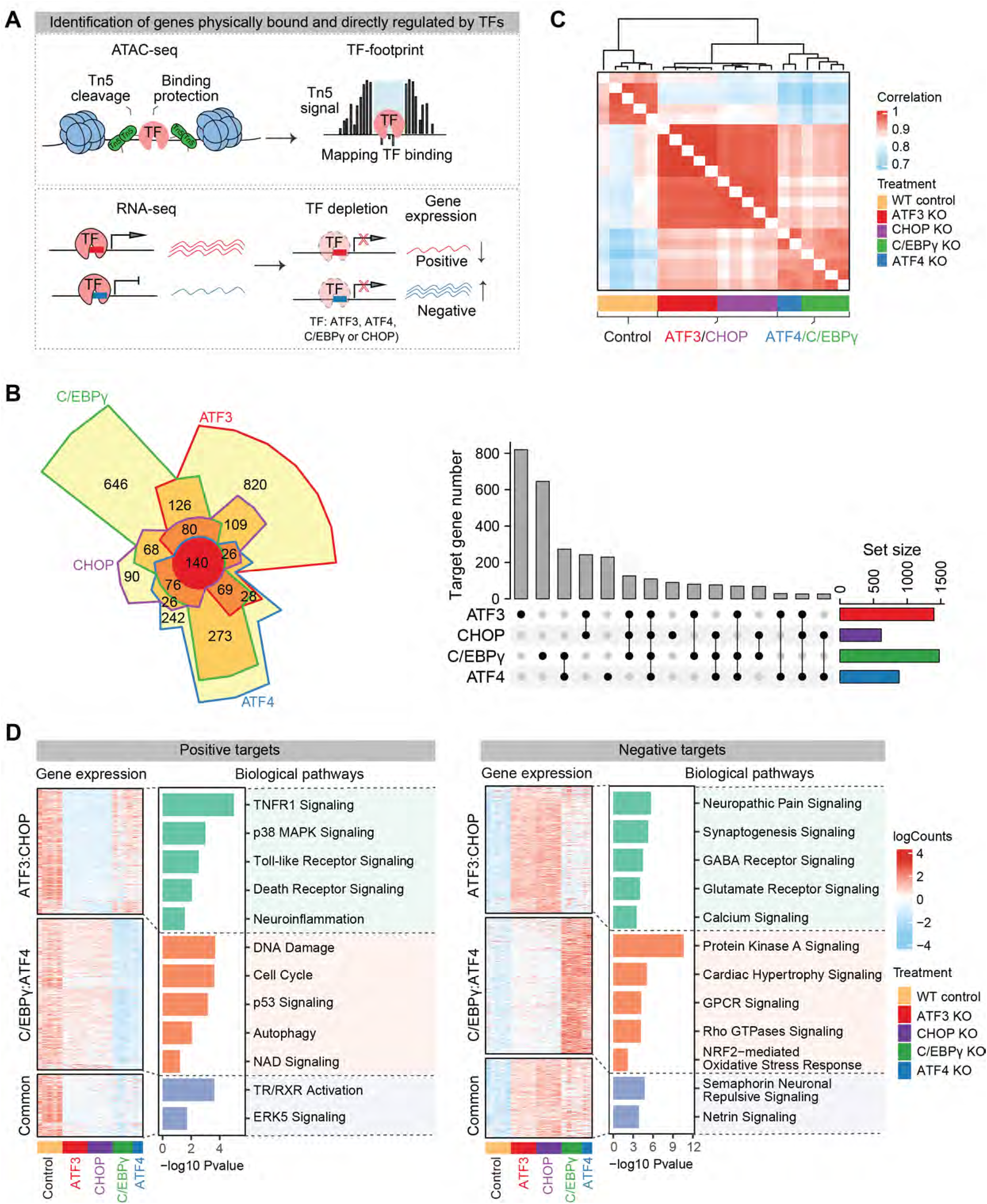
Identification of direct target genes of the four survival TFs. (A) A schematic diagram displaying integrative analysis of DNA-footprinting using ATAC- seq data and RNA-seq to identify each TF’s direct target genes. Chromatin-bound TF protects DNA elements from Tn5 transposase cleavage, creating single-nucleotide- resolution DNA “footprints” during ATAC-seq. Mapping these footprints would identify DNA regions directly bound by the TF, and the genes linked to TF footprinted regions. To identify genes that are directly regulated by each survival TF, RNA-seq was performed on RGCs with or without CRSIPR ablation of this TF at day 3 following injury. Genes that are footprinted by this TF and differentially expressed (up- or down-regulated, absolute logFC >0 and FDR < 0.1) upon ablation of this TF are considered as its direct target genes. (B) A Chow-Rusky Venn diagram and an Upset plot showing the overlap of each TF’s direct targets. The set size indicates the number of direct targets for each TF. The intersection size indicates the number of overlapped targets unique to dotted TF(s). (C) A heatmap of correlation matrix showing similarity and dis-similarity in the expression levels of each TF’s direct target genes. Normalized RNA-seq counts of each TF’s direct target genes were correlated and clustered by the distances among samples from control RGCs (nontargeting sgRNA) or treated RGCs receiving individual CRISPR TF ablations. Colors in the heatmap indicate Pearson’s correlations between two samples. ATF3 and CHOP are more similar in the expression levels of their target genes, while ATF4 and C/EBPγ are more similar. (D) Shared or unique target gene sets of each TF. Positive targets are defined as genes that are down-regulated (log2FC < 0, FDR < 0.1) when this TF is ablated. Negative targets are defined as genes that are up-regulated (log2FC > 0, FDR < 0.1) when this TF is ablated. Based on gene expression similarity in (C), each TF’s positive or negative targets were grouped together and clustered into three gene sets: a) genes that are uniquely controlled by ATF3 and CHOP, b) genes that are uniquely controlled by ATF4 and C/EBPγ, and c) genes that are shared by ATF3/CHOP and ATF4/C/EBPγ groups. Gene ontology on these three gene sets were performed and related GO terms with enrichment FDR value < 0.05 were presented.

The second method to characterize downstream, survival regulating pathways was to perform RNA-seq on injured RGCs from which ATF3, ATF4, C/EBPγ or CHOP had been deleted (Figure 4A). We injected AAVs expressing gene-specific sgRNAs and mCherry reporter to the vitreous bodies of LSL-Cas9: Vlgut2-Cre mice, so that sgRNA-mediated knockouts of individual TFs occur selectively in RGCs (Zhang et al., 2019). At day 3 following optic nerve crush, transduced RGCs were FACS-purified for RNA-seq. Following quality control and outlier removal, we analyzed gene expression changes caused by each perturbation. We found that in comparison to control (non-targeting) gRNAs, knocking out ATF3, ATF4 or CHOP resulted in ∼1,800 differentially expressed genes (DEGs), whereas C/EBPγ depletion generated ∼5,600 DEGs (Figure S4A, Table S5). Recognizing that genes identified by RNA-seq include both direct and indirect targets of specific TFs, we focused on the overlap between each TF’s DEGs from RNA-seq coupled to TF perturbations and its target genes predicted by footprint analysis (Table S6). We observed significant overlaps, with ∼27-50% of the knockout-associated DEGs, identified as direct target genes of individual TFs (Figure S4B), supporting our hypothesis that RNA-seq identifies both direct and indirect transcriptional responses.

### Identification of Two Degenerative Programs Dependent on ATF3/CHOP and C/EBPγ/ATF4 Respectively

Further analyses focused on the direct transcriptional targets of ATF3, ATF4, CHOP and C/EBPγ. As expected, direct targets include both down-regulated or up-regulated genes due to CRISPR knockout of each TF (Figure S4C), thus representing positively or negatively regulated genes of the TFs, respectively. By comparing the GO terms of these target genes of individual TFs, we found that the shared functions of these TFs are relevant to regulating responses to stimulus and cell death processes (Figure S4D), consistent with their roles in mediating RGC degeneration after injury (Figure S4D). Interestingly, several of these target genes, such as Srebf1 and Rora, which are shared targets of all 4 TFs, have been implicated in human glaucoma by a GWAS study (Gharahkhani et al., 2021).

To assess overlap among gene expression programs regulated by the four key TFs, we constructed a Venn diagram of their direct target genes. As shown in Figure 4B, ATF3 and C/EBPγ each regulate a unique set of genes, while most target genes of ATF4 or CHOP are shared by the other two TFs. Hierarchical clustering on the expression levels of the direct target genes of individual TFs revealed distinct gene expression patterns in injured RGCs after perturbation, one shared by knockout of ATF3 or CHOP, and the other by the knockout of ATF4 or CEBPγ (Figure 4C). We therefore group these directly targeted genes and their GO terms into different clusters: positive or negative targets shared by ATF3/CHOP, C/EBPγ/ATF4, or those shared by all four TFs (Figure 4D).

Strikingly, ATF3/CHOP and ATF4/CEBPγ appear to regulate different positive and negative regulatory pathways (Figure 4D and 5A). The positive targets of ATF/CHOP mainly regulate neuronal responses to extrinsic factors, such as TNFR1/p38 MAPK signaling, Toll-like receptor signaling and neuroinflammation. In contrast, ATF4/C/EBPγ’s positive targets are more relevant to cell intrinsic stress responses, such as cell cycle/DNA damage/checkpoint regulation, p53 and NAD signaling and autophagy. For example, Tnfrsf21 or death receptor 6 (DR6), a TNF receptor family member that triggers cell death (Benschop et al., 2009), is uniquely bound and activated by ATF3 and CHOP, but not C/EBPγ and ATF4 (Figure 5A-B, ATF3/CHOP positive; Figure S5). In contrast, Hdac9, which regulates endogenous DNA repair (Wong et al., 2009), is a unique target of C/EBPγ and ATF4 (Figure 5A-B, C/EBPγ/ATF4 positive; Figure S5). Negative targets of ATF3/CHOP and ATF4/CEBPγ are also distinct. ATF3/CHOP negative targets are mainly associated with different aspects of synaptic function, suggesting that ATF3 and CHOP mediate direct injury-triggered disruption of RGCs’ physiology. In contrast, the negative targets of ATF4/C/EBPγ are highly relevant to protein kinase A signaling and Rho GTPase regulation, important processes related to intrinsic regulation of signaling and homeostasis. Thus, these results revealed two complementary programs in mediating injury-induced RGC degeneration.

**Figure 5.**
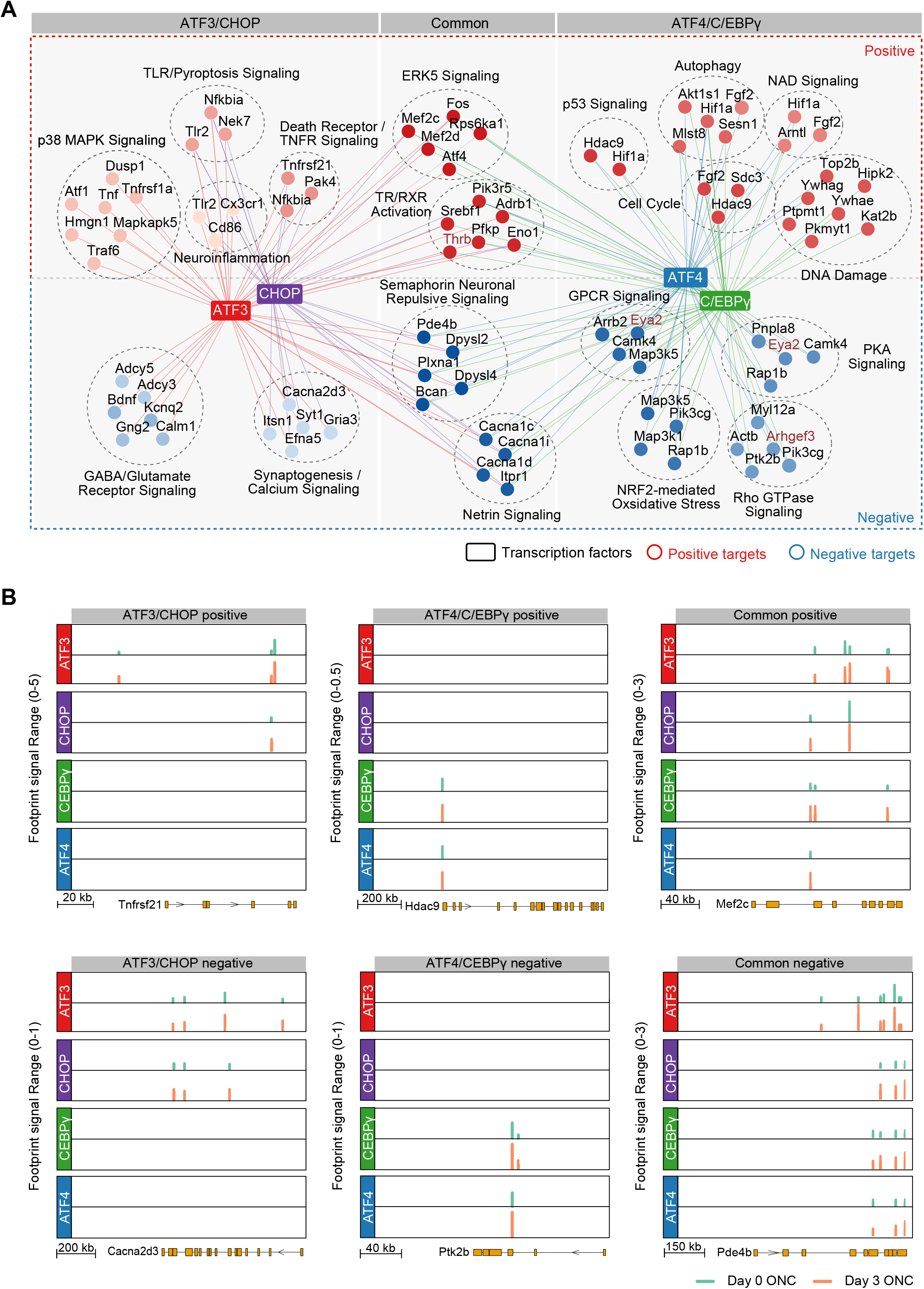
Two distinct transcriptional programs regulated by ATF3/CHOP or C/EBPγ/ATF4 in injured RGCs. (A) A TF network plot showing the positive or negative targets and the associated biological pathways that are uniquely controlled by ATF3 and CHOP (ATF3/CHOP), uniquely controlled by ATF4 and C/EBPγ (ATF4/C/EBPγ), or shared by both ATF3/CHOP and ATF4/C/EBPγ (common). Representative genes in each pathway were shown. Each node indicates a gene. Node colors indicate the different GO terms, and edge colors indicate TFs. (B-C) Genome browser views of each of 4 TF’s binding sites around representative target genes. Y-axis indicates Tn5-bias corrected footprint signals of each TF from control (day 0) or optic nerve crushed (day 3) RGCs. Shown representative genes, including both positive (B) and negative (C) targets, are selected from the three groups in (A).

### Additive Effects of Co-knockout of TFs in Both Divisions Resulting in nearly Complete Protection of RGCs After Injury

If ATF3/CHOP and CEBPγ/ATF4 indeed regulate distinct degenerative pathways in injured RGCs, co-manipulating one TF from each of the two subgroups would be predicted to have additive effects and further improve RGC survival compared to individual CRISPR perturbations of each TF. To test this prediction, we deleted several pairs of the four TFs and assessed neuronal survival. Injury-induced alterations could regulate RGC survival by RGC- autonomous and RGC- nonautonomous mechanisms (Zhang et al., 2019). We revised our protocol by injecting gene-specific sgRNA-carrying AAVs to the vitreous bodies of the mice of LSL-Cas9: Vglut2-Cre. In this way, sgRNA-mediated knockouts of individual TFs occur selectively in RGCs but not in other retinal cell types (Zhang et al., 2019)(Figure S6A). As shown in Figure S6B and S6C, RGC-selective knockout of each of these TFs led to virtually identical survival outcomes in these mice as non-selective knockouts (Figure 1C and 1D), supporting the RGC-autonomous effects of these TFs.

To minimize the effect of diluting AAV2 vectors due to combining different pools of sgRNAs, we intravitreally injected two different sgRNA encoding AAV2 vectors together with AAV2-Cre into LSL-Cas9 (Figure 6A) and verified knockout efficiency by immunohistochemistry (Figure 6B-E, S6D-G). Co-ablation of one TF from each of the two subgroups led to higher RGC survival in three of four cases (ATF3/CEBPγ, ATF3/ATF4, and CEBPγ/CHOP) compared to single knockout (Figure 6F, 6G). In contrast, co-knockout of TFs within the same group, (ATF3/CHOP or CEBPγ/ATF4) was not significantly more protective than single knockout (Figure 6F, 6G). Considering that 80-90% RGCs are transduced by intravitreal injected AAVs (Jacobi et al., accompany paper), these results suggested that co-knockout of these TFs protect most AAV-transduced RGCs from injury-induced degeneration. Thus, these observations provide functional validation of our bioinformatic predictions from integrative analysis of DNA-footprinting and RNA-seq, supporting the overlapping, yet complementary effects of ATF3/CHOP and CEBPγ/ATF4 on activating two parallel pro-death pathways following injury.

**Figure 6.**
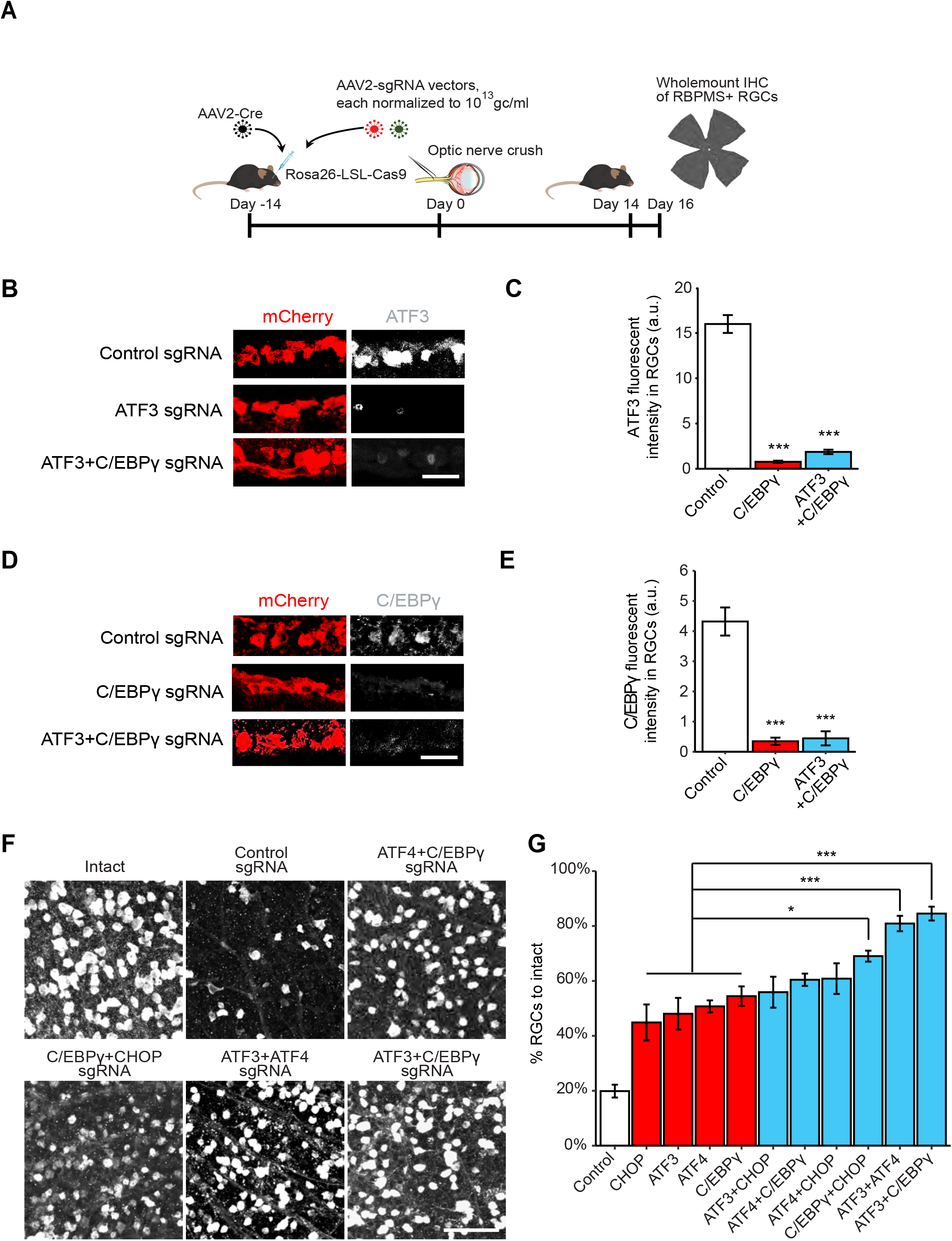
Functional interactions of the key TFs in injured RGCs. (A) Schematic illustration of CRISPR KO with sgRNAs targeting multiple genes. (B-C) Representative images (B) and quantification (C) of ATF3 immunohistochemical staining indicate the knockout efficiency of ATF3 sgRNA and ATF3+C/EBPγ sgRNA. Data are shown as mean ± s.e.m. with n = 4. ***p<0.001. Scale bar, 10 μm. (D-E) Representative images (D) and quantification (E) of C/EBPγ immunohistochemical staining indicate the knockout efficiency of C/EBPγ sgRNA and ATF3+C/EBPγ sgRNA. Data are shown as mean ± s.e.m. with n = 4. ***p<0.001. Scale bar, 10 μm. (F) Representative images of RGC survival with combinations of sgRNA hits injected to LSL-Cas9 mice. Scale bar, 50 μm. (G) Quantification of RGC survival with combinations of sgRNA hits. Data are shown as mean ± s.e.m. with n = 4-5 biological repeats. *p<0.05, ***p<0.001.

### Knockout of Identified TFs Protects RGCs in a Glaucoma Model

Identification of these transcriptional drivers of injury-induced neuronal degeneration and their downstream pathways provides new potential therapeutic targets. To directly test whether our findings extend to pathological conditions, we extended our knockout analysis to a glaucoma model. Glaucoma, a leading cause of irreversible blindness worldwide, is the result of progressive RGC loss (Quigley and Broman, 2006; Weinreb et al., 2014). Elevated intraocular pressure (IOP) is a major risk factor, and a proposed pathological mechanism is initial axonal damage and subsequent RGC death (Quigley et al., 1983; Sommer et al., 1989; Calkins et al., 2012; Nickells et al., 2012; Weinreb et al., 2016; Guo et al., 2021). Likewise, ocular hypertension in rodents leads to RGC death and has been widely used to model glaucoma (Sappington et al., 2010, Cone et al., 2010; Zhang et al., 2019). To induce consistent IOP elevation, we modified the conventional microbead/viscoelastic strategy and developed a novel viscobead occlusion experimental glaucoma model (Figure S7 and 7A-G). After injection into the anterior chamber, the viscobeads rapidly accumulated at the iridocorneal angle (Figure S7E-F, 7A-C) and induced consistent IOP elevation for at least 8 weeks in most mice (Figure 7D and S7F). Importantly, about 30% of RGC loss was consistently observed at 8 weeks after viscobeads injection, as assessed in retinas (Figure 7E-G) and optic nerves (Figure S7G-K).

**Figure 7.**
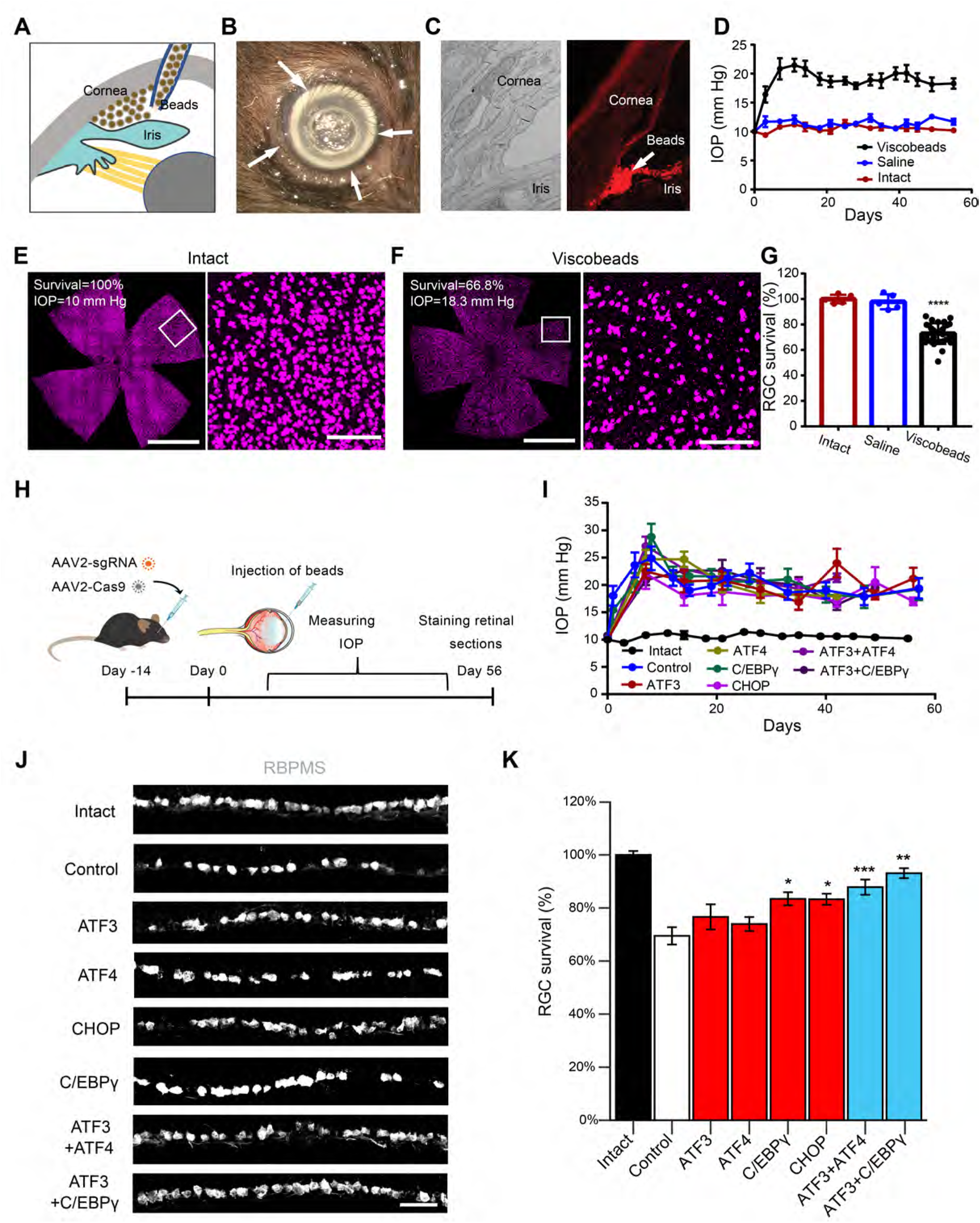
CRISPR ablation of the identified TFs protects RGCs in a glaucoma model. (A) Schematic illustration of the viscobeads occlusion experimental glaucoma model. Concentrated viscobeads were injected into the mouse anterior chamber to block aqueous drainage. (B) A representative photograph of viscobeads accumulated at mouse iridocorneal angle 5 min after injection. Arrows show the white viscobeads were restricted at the iridocorneal angle. (C) Viscobeads distribution in the mouse anterior chamber following intracameral injection. Left: A representative transmission electron microscopy (TEM) image of an intact mouse iridocorneal angle. Right: A representative fluorescence image showing rhodamine B labelled viscobeads accumulation at mouse iridocorneal angle. Scale bar, 10 μm (left) and 100 μm (right). (D) Intraocular pressure (IOP) in the mice before and after the injection of viscobeads (n=10) or saline (n=5) and the naïve group (n=5). Data are shown as mean ± SD. (E, F) Representative wholemount retina confocal images from an intact mouse (E) and a mouse after 8 weeks from viscobeads injection (F). RBMPS antibody was used for the immunohistochemical staining of RGCs. Scale bar for the left panel for (F, G), 2 mm. Scale bar for the left panel for (F, G), 100 μm. (G) Quantification of RGC survival 8 weeks of after the viscobeads (n=31) or saline (n=5) injection and the naïve group (n=5). Data are shown as mean ± SD. (H-K) Knockout of individual TFs protects RGCs in the glaucoma mice. (H) Schematic illustration. (I) IOP elevation in different group of mice. (J) Representative images of retinal sections showing RGC survival with sgRNA injections targeting indicated TFs. Scale bar, 20 μm. (K) Quantification of RGC survival with sgRNA injections targeting the key TFs. Data are shown as mean ± s.e.m.

To ask whether the knockdown of ATF3, ATF4, CEBPγ or CHOP, alone or in combination, would increase RGC survival, we injected the AAV2 vectors encoding the single-targeting or multiple-targeting sgRNAs together with AAV2-Cas9 (Wang et al., 2020) into the vitreous bodies of wildtype mice that had received bead injections two weeks previously (Figure 7H). None of the AAV injections altered IOP elevation (Figure 7I). However, in comparison to control sgRNA group, knockout of C/EBPγ and CHOP showed a statistically significant increase in RGC survival under the same level of elevated IOP (Figure 7J-K). Furthermore, we observed additive RGC protection effects for both ATF3+ATF4 and ATF3+C/EBPγ CRISPR groups, similar to our observations in the ONC model (Figure 6F-G). These results support the transability of these TFs and their target downstream pathways in multiple contexts. The TFs defined here may therefore represent synergistic molecular targets for neuroprotection after injury.

## DISCUSSION

Through two orthogonal genome-wide strategies, *in vivo* CRISPR screening and combined assays of chromatin accessibility (ATAC-seq) and gene expression (RNA-seq), we identified several TFs that play key roles in injury-induced neuronal loss. We then focused on a set of four TFs – ATF3, ATF4, CEBPγ and CHOP identified through these independent approaches. We showed that their activation and complementary action play key roles in the early engagement of different degeneration-regulatory pathways in injured RGCs. Manipulation of these TFs also led to significant neuroprotection in a newly modified glaucoma model.

Previously, several hypotheses have been proposed to explain axotomy-induced neuronal degeneration. For example, axotomy may deprive target-derived pro-survival signals, or initiate the production of “injury signals” that can be retrogradely transported to the cell bodies to elicit degenerative responses (Howell et al., 2013; Syc-Mazurek and Libby, 2019). In addition, injury may trigger rapid responses in surrounding cells, including connected neurons and glial cells along the nerve and in the retina, thus indirectly triggering neuronal responses (Liddelow et al., 2017, Baldwin et al., 2015; Williams et al., 2020). Therefore, it was thought that there would be many TFs and pathways regulating injury-elicited neuronal survival/death. Surprisingly, our large-scale CRISPR screening identified only a small number (10 out of 1893) of TFs that negatively regulate neuronal survival, which remarkably, are concentrated in the family of basic leucine zipper domain (bZIP), in particular, ATF and C/EBP members. Furthermore, four of these positive hits (ATF3/4, CEBPγ and CHOP) were also identified as key injury regulators by independent studies of chromatin accessibility coupled to TF footprinting, which prompted us to focus on these candidate master regulatory genes. Our results showed that knockout of each of these four TFs individually significantly improves neuronal survival, while knockout of specific combinations had additive effects, which were remarkably consistent with our bioinformatic analyses of their distinct and overlapping regulatory targets. Furthermore, expression of these TFs is reduced in injured RGCs with pro-survival interventions, including Pten/Socs3 deletion with or without CNTF (Jacobi et al., accompanying paper). Encouraging protective effects of manipulating these TFs in an experimental glaucoma model points to these TFs as generalizable therapeutic targets to prevent neuronal loss.

How might these TFs act in injured RGCs? Our results suggest that they are activated (both by expression and binding activity) at very early time points post-injury. Important mechanistic insights came from our analysis of functionally relevant direct target of these TFs, as activators or repressors. As expected, some targets shared by the four TFs are functionally relevant to cell death, such as ERK5 signaling, which has been previously implicated in mediating neurotrophin-mediated retrograde survival responses (Watson et al., 2001; Heerssen and Segal, 2002). Remarkably, our results revealed that ATF3/CHOP and CEBPγ/ATF4 engage in two distinct neurodegeneration programs. ATF3/CHOP preferentially up-regulate genes related to neuronal responses to extrinsic factors, such as TNFR/IRF and neuroinflammation, but down- regulates genes related different aspects of synaptic transmission function. In contrast, genes induced by CEBPγ/ATF4 are more relevant to cellular responses to intrinsically generated stressors, such as the DNA damage response and the NAD/p53 pathway, perhaps relating to injury-triggered metabolic reprogramming and oxidative stress in injured RGCs. This might help mechanistically explain the results of a recent study showed that a sciatic nerve injury induces double-stranded breaks in axotomized sensory neurons (Cheng et a., 2021). Furthermore, CEBPγ/ATF4-suppressed genes are highly relevant to the intrinsic homeostatic processes, such as PKA and Rho signaling, which are crucial for neuronal survival regulation (Goldberg, 2004; Huang and Reichardt, 2001). Interestingly, several of the genes regulated by TFs have been implicated in human glaucoma by GWAS studies (Gharahkhani et al., 2021). Future studies will analyze the mechanistic basis of these TFs in regulating these diverse transcriptional programs.

In addition to RGC degeneration, some of these molecules and pathways have been suggested as important players in other types of neurodegeneration. For example, neuronal innate immune activation, a targeted process of ATF3/CHOP shown here and manifest by neurons expressing mutant Huntingtin (Lee et al., 2020) and p53, is also a key regulator of c9orf72 poly(PR) triggered neurodegeneration (Maor-Nof et al., 2021). However, considering our finding that a seemingly simple axotomy could rapidly elicit multiple degenerative cascades in RGCs, it is conceivable that these different degeneration programs might be also co-activated in other neurodegenerative diseases, which might contribute to largely unsatisfactory results from the current state of the field that relies only on single-targeted treatment of neurodegenerative diseases. Thus, our results highlight the potential importance of manipulating multiple targets for effective neuroprotection.

Our analysis of the four TFs also revealed important differences between responses of CNS and PNS neurons to injury. For example, axotomized primary sensory neurons are known to have marked ATF3 up-regulation but do not die; instead, they mount regenerative responses (Renthal et al., 2020; Cheng et al., 2021). Thus, ATF3 is a critical pro-death gene in RGCs, but not in peripheral sensory neurons. Furthermore, whereas ATF4/CEBPγ/CHOP are upregulated in injured RGCs, they exhibit minimal or no up-regulation in injured sensory neurons (Renthal et al., 2020). It is also interesting to note that although neuronal death is the most prominent pathological event in CNS neurodegenerative diseases, it rarely occurs in peripheral neuropathies, consistent with our findings. Thus, it would be important for future studies to explore the mechanisms that accounts for different expression and action of these TFs in the PNS and CNS, which may provide crucial understanding on the differences in cell death and subsequent repair observed after injury.

A prerequisite of axon regeneration is neuronal survival. But it is less known how these processes are coordinated in injured neurons. Our results provide several insights. First, the CRISPR screen identified largely separate sets of TFs negatively regulating survival and regeneration. While most survival genes are ATF/CEBP members, most identified regeneration-inhibiting TFs are implicated in epigenetic control of gene expression. Furthermore, knockout of these epigenetic factors fails to promote neuronal survival, and their regeneration phenotypes are weaker than what is seen after knockout of PTEN and/or SOCS3 (Park et al., 2008; Smith et al., 2009; Sun et al., 2011; Bei et a., 2006). These results suggest that such epigenetic mechanisms may provide permissive control of axon regeneration, different from instructive regulation provided by PI3K/mTOR and JAK/STAT activation consequent of PTEN or SOCS3 inhibition (Smith et al., 2009; Sun et al., 2011; Jacobi et al., accompanying paper). Second, our results reveal both distinct and common effects of TFs on axon regeneration in CNS and PNS. For example, CTCF knockout inhibits axon regeneration in PNS (Palmisano et al., 2019), in stark contrast to its regeneration-promoting effects of its knockout in ONC, as shown here (Figure 1F). On the other hand, ATF3 is upregulated in both DRG and RGCs after injury, and it acts as a positive regulator of axon regeneration in both cases (Renthal et al., 2020; Kole et al., 2020). Indeed, ATF3 knockout completely abolished the regeneration phenotype triggered by PTEN deletion (Jacobi et al., accompanying paper). In light of the massive numbers of direct and indirect targets of ATF3 (this study, Renthal et al., 2020), we speculate that ATF3 may act as reprogramming factor, permitting the activation of axon regeneration program in injured neurons, even though such reprogramming process may render neurons more prone to cell death. Future studies are needed to develop combinatorial treatments that not only protect injured neurons from degeneration, but also promote their regenerative programs.

## STAR★METHODS

### KEY RESOURCE TABLE

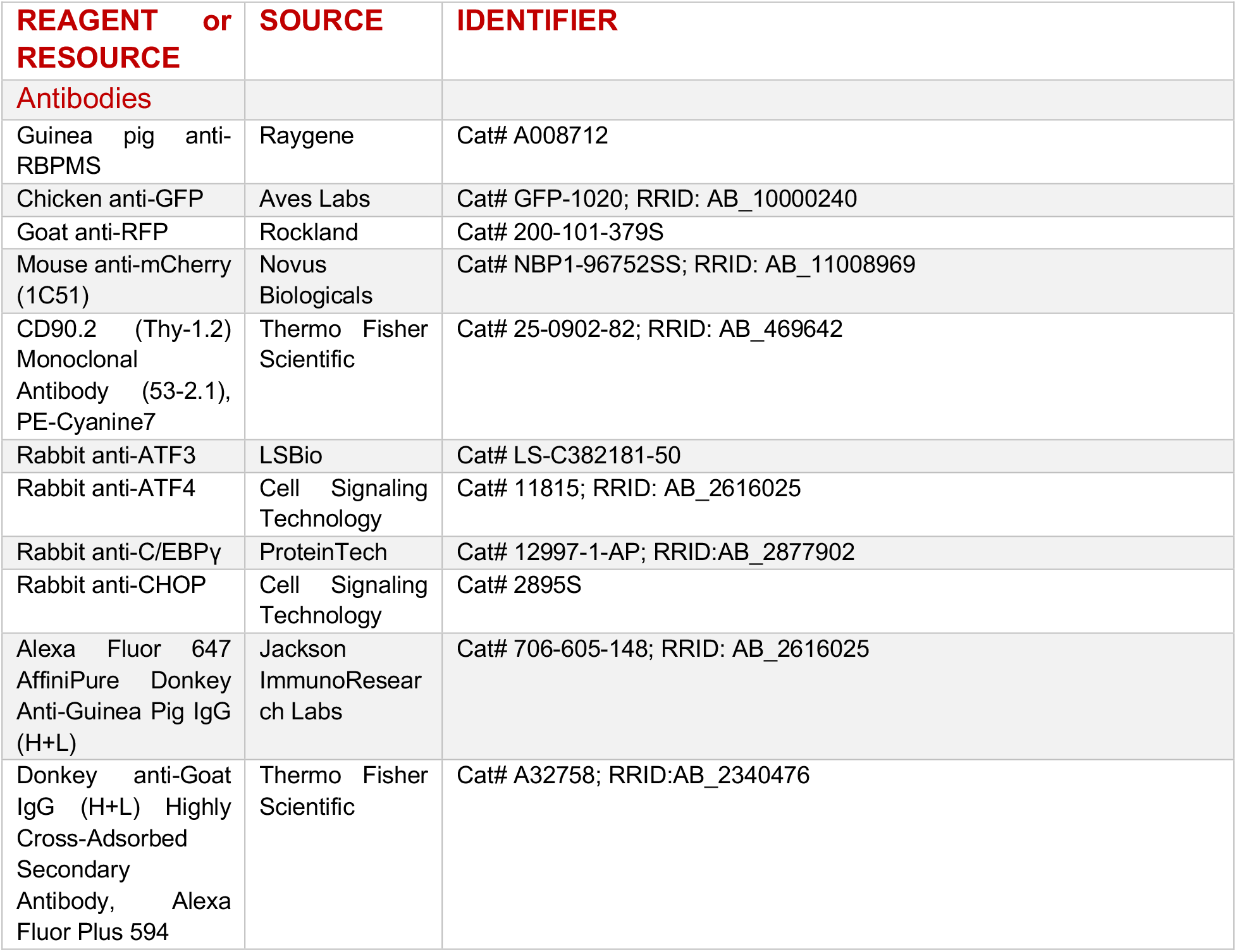
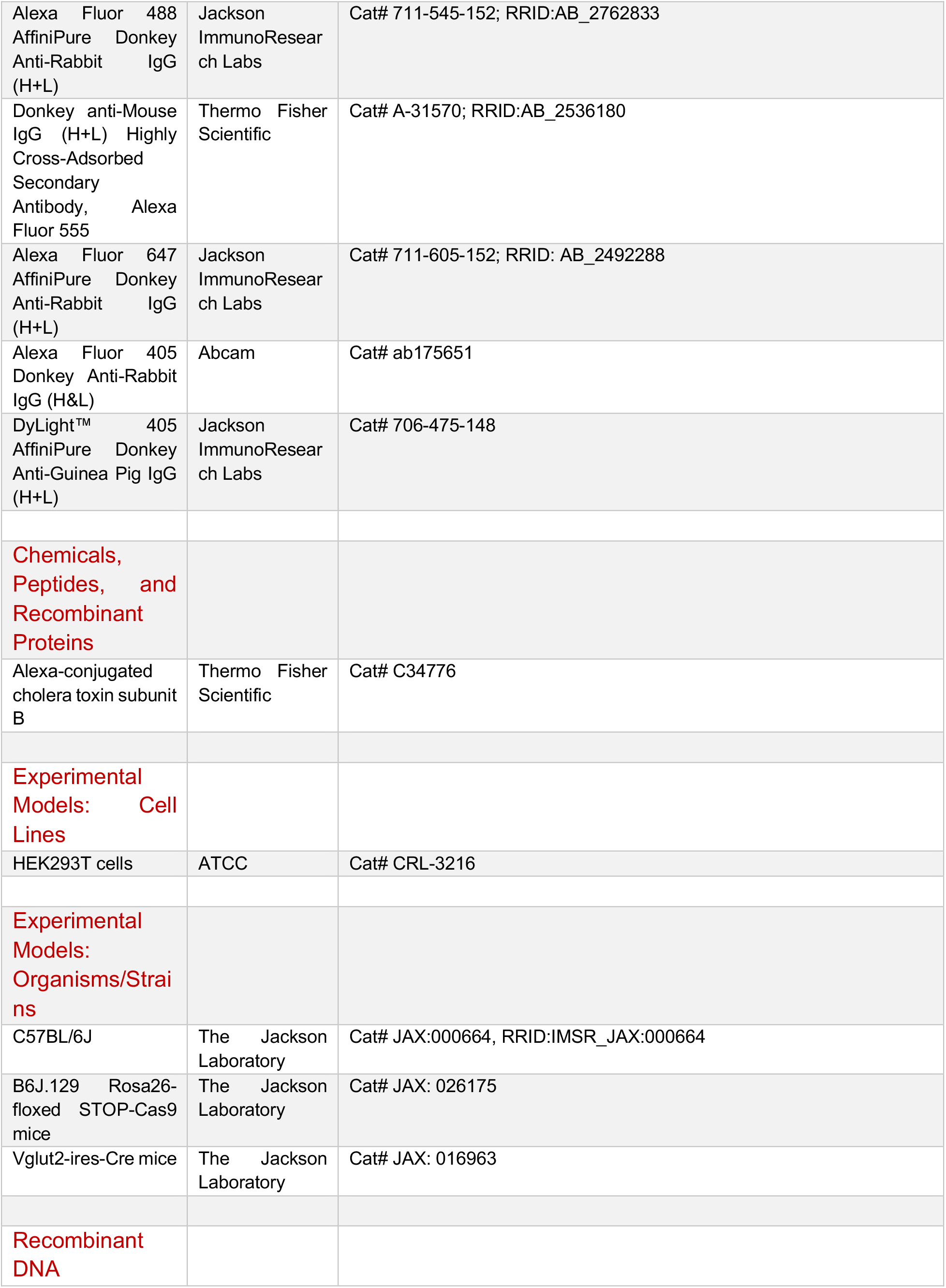
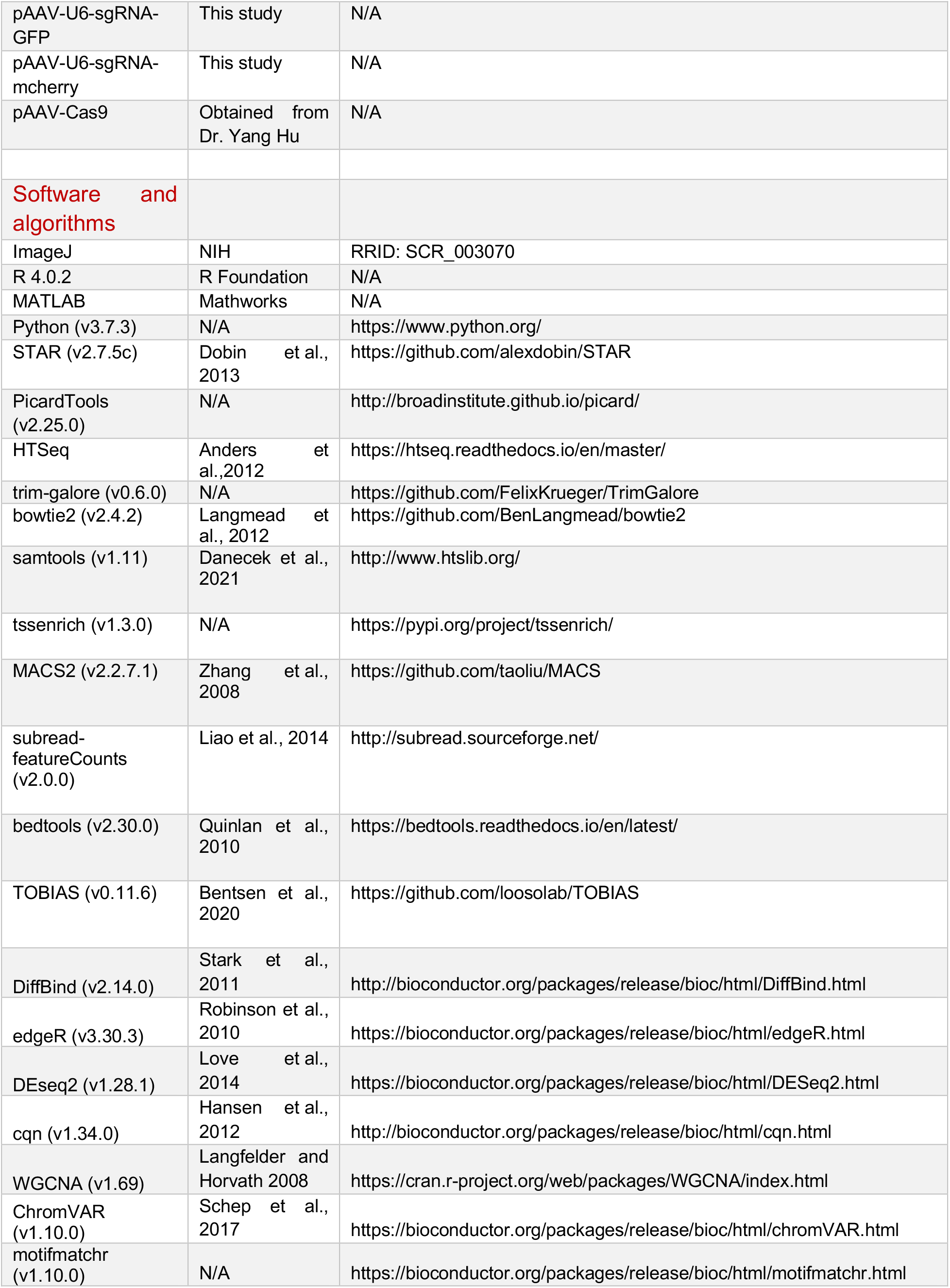
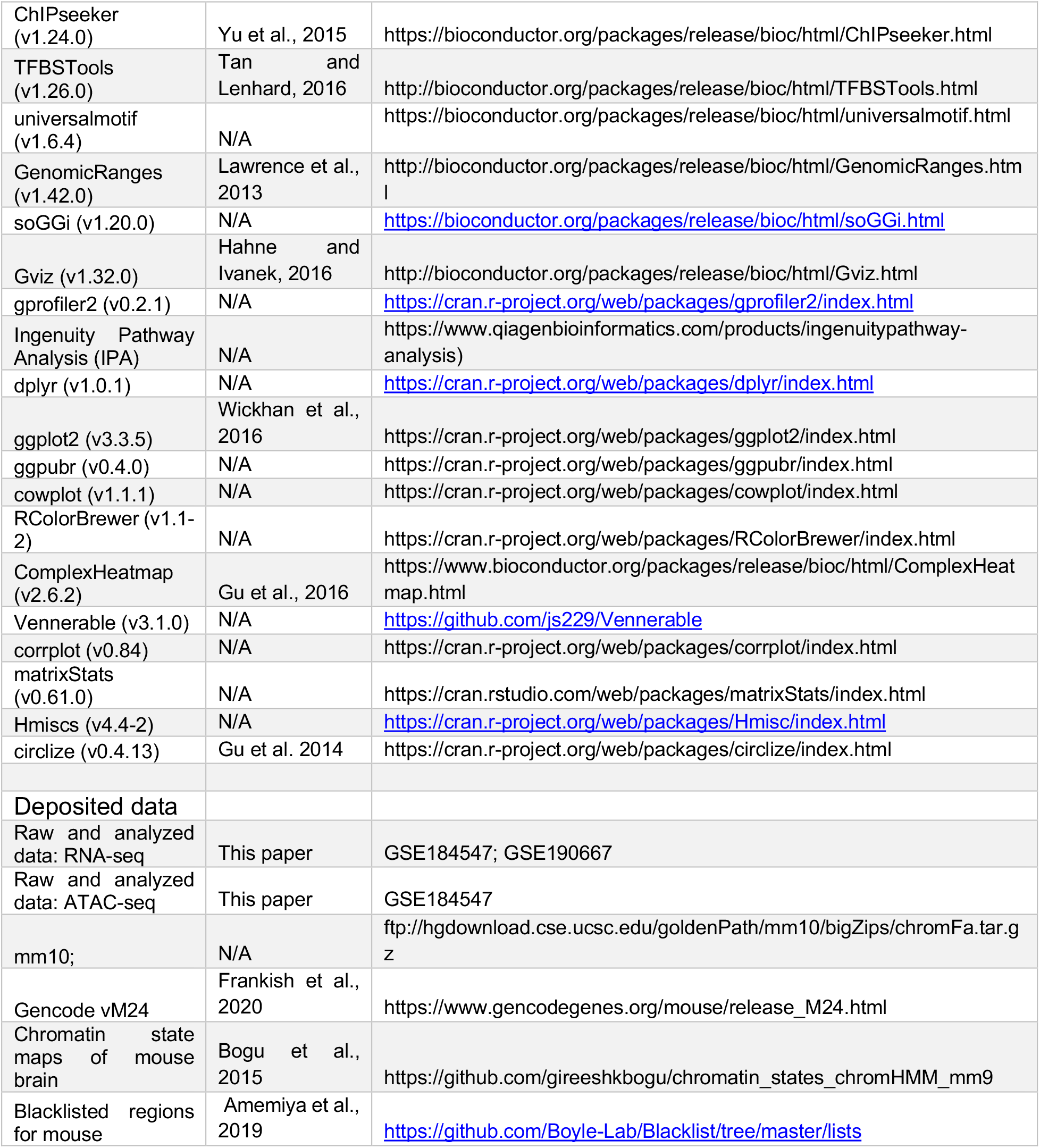

### EXPERIMENTAL MODEL ANDRESOURCE AVAILABILITY

### EXPERIMENTAL MODEL AND SUBJECT DETAILS

#### Animals

All experimental procedures were performed in compliance with animal protocols approved by the IACUC at Boston Children’s Hospital and Harvard University. Mice aged at 4 weeks were used for optic nerve crush model. Male and female mice were used in this study at ratios dependent on litters available and with equal distributions across experiments conducted extemporaneously. B6J.129 Rosa26-floxed STOP-Cas9 (Rosa26-LSL-Cas9) (026175; Jackson Labs) and Vglut2- ires-Cre mouse strains (016963; Jackson Labs) were obtained from Jackson Laboratories. The ATF3^f/f^ mouse strain was acquired from Dr. Clifford Woolf’s lab (Renthal et al., 2020).

### METHOD DETAILS

#### Library Virus production

First, we utilized the RIKEN databases (Kanamori et al., 2004; Fulton et al., 2009) and generated a non-redundant list of 1893 transcription regulators across nearly the entire mouse genome. Second, to maximize knockout efficiency, we selected five sgRNAs targeting different regions of each TF from the well-characterized genome-scale CRISPR/Cas9 knockout (GeCKO) library (Shalem et al., 2014; Shalem et al., 2015). These sgRNAs were subcloned into pAV-U6-gRNA (Swiech et al., 2015) and the resultant expression vectors with sgRNAs for the same TF gene were mixed. As a result, we generated a library of 1893 pooled sgRNAs, each of which has 5 sgRNA-bearing plasmids for individual TF genes. Third, these plasmids were used to prepare AAV vectors for transducing RGCs in vivo. We chose serotype 2/2 (AAV2) because it transduces RGCs following intravitreal injection with high efficiency and reasonable selectivity (mostly RGCs and amacrine cells in the ganglion cell layer) (Nawabi et al., 2015; Norsworthy et al., 2017; Park et al., 2008). All AAV viral vectors were made by Boston Children’s Hospital Viral Core. AAV serotype 2 were used in our study as following: AAV2-Cre; AAV2-PTEN sgRNA; AAV2-sgRNAs for other screened TFs. The titers of all viral preparations were at least 5 x 10^12^ genome copies/mL for sgRNA injection targeting single gene. Viral titers for sgRNA combinations targeting multiple genes were adjusted to 1 x 10^13^ genome copies/mL for each target gene.

#### Surgical procedures

For all surgical procedures, mice were anaesthetized with ketamine and xylazine and received Buprenorphine as a postoperative analgesic.

##### Intravitreous AAV injection

As previously described, intravitreal virus injection was performed two weeks before optic nerve crush injury to enable axon regeneration. Briefly, a pulled-glass micropipette was inserted near peripheral retina behind the ora serrata and deliberately angled to avoid damage to the lens. 2 ul of the combination of AAV2/2-CAG-Cre virus and AAV2-sgRNAs were mixed (1:3 mix) was injected for LSL-Cas9 mice (Platt et al., 2014). 2 ul AAV2/2 virus was injected for or LSL-Cas9; vGLUT2-Cre mice. For sgRNA combinations targeting multiple genes, the titer of each sgRNA encoding AAV was adjusted to 1 x 10^13^ genome copies/mL and mixed at a ratio of 1:1.

##### Optic nerve injury

As previously described, optic nerve was exposed intraorbitally and crushed with fine forceps (Dumont #5 FST) for 5s, approximately 500um behind the optic disc. Afterwards, eye ointment was applied post-operatively to protect the cornea. Robust axon regeneration could be observed from 2 weeks post crush by Alexa-conjugated cholera toxin subunit B labeling.

#### Perfusions and tissue processing

For immunostaining, animals were given an overdose of anesthesia and transcardiacally perfused with ice cold PBS followed by 4% paraformaldehyde (PFA, sigma). After perfusion, optic nerves were dissected out and postfixed in 4% PFA overnight at 4⁰C. Tissues were cryoprotected by sinking in 30% sucrose in PBS for 48 hours. Samples were frozen in Optimal Cutting Temperature compound (Tissue Tek) using a dry ice and then sectioned at 12 mm for optic nerves.

#### Retinal wholemount staining and quantification of RGC survival

Dissected retinas were rinsed in PBS and then blocked in PBS with 1% Triton X-100 and 5% horse serum (wholemount buffer) overnight at 4 °C. Primary antibodies diluted in the wholemount buffer were then treated for 2-4 days at 4 °C, followed by three times of rinsing by PBS (10 min each time). Secondary antibodies (all with 1:500 dilution) were diluted in PBS and treated overnight at 4 °C. After five times of rinsing by PBS (10 min each time), retinas were mounted with Fluoromount-G (Southern Biotech, Cat. No. 0100-01).

#### Immunostaining and imaging analysis

Cryosections (12-μm thick) were permeabilized and blocked in blocking buffer (0.5% Triton X-100 and 5% horse serum in PBS) for 1 h at room temperature and overlaid with primary antibodies overnight at 4 Celsius degree (Table 1). For BrdU staining, cells or tissue sections were denatured with 2 N HCl for 30 minutes at 37 Celsius degree and then were neutralized with 0.1 M sodium borate buffer for 10 min before proceeding to normal blocking procedure. On the next day, the corresponding Alexa Fluor 488-, 594- or 647-conjugated secondary antibodies were applied afterwards (all secondary antibodies were purchased from Invitrogen). All stained sections were mounted with solutions with DAPI-containing mounting solution and sealed with glass coverslips. All immunofluorescence-labeled images were acquired using Zeiss 700 or Zeiss 710 confocal microscope. For each biological sample, 3-5 sections of each optic nerve were imaged were taken under 10x or 20x objectives for quantification. Positive cell numbers were then quantified manually using the Plugins/ Analyze /Cell Counter function in ImageJ software. For fluorescent intensity analysis, the images were first converted to 8-bit depth in ImageJ software and then, the mean intensity value was calculated by the build-in function: Analyze/Measure.

#### Tissue clearing, imaging and quantification of optic nerve regeneration

Mice injected with fluorophore tagged Cholera Toxin B (CTB) were perfused with 4% paraformaldehyde. Dissected optic nerves were then subjected to a modified procedure from previously published iDISCO tissue clearing method, which rendered the optic nerves transparency for direct fluorescent imaging (Renier et al., 2014). This procedure has been tested for better preservation of CTB florescence and the least change of optic nerve shape during tissue clearing. For dehydration, optic nerve samples were incubated in dark for 0.5h of 80% tetrahydrofuran (THF, Sigma-Aldrich 360589-500ML)/H_2_O and then switched to 100% THF for 1h. Then, samples were incubated in Dichloromethane (DCM, Sigma-Aldrich 270997-1L) for 20min (nerves should sink at the bottom). Samples were finally switched to dibenzyl ether (DBE, Sigma-Aldrich 33630-250ML) until complete transparency (at least 3h, but overnight is recommended). Transparent nerves can be stored in DBE without obvious fluorescence decay of CTB for at least 1 year. For imaging, processed nerves can be mounted in DBE and imaged under Zeiss 710 confocal microscope. Z-stack scanning and maximum projection of Z-stack images were used in order to capture all regenerated axons. For image analysis, fluorescent intensity profile along the nerve was generated by the build-in function of ImageJ: Analyze/Plot Profile. The crush site of maximum projected optic nerve image was identified by overlaying 5 single slice images at different imaging depth (z-direction) of the same nerve (Figure S1A). To calculate the integral of fluorescent intensity across the entire length of the nerve, a Matlab algorithm was developed by our lab to quantify the “area under curve” from the plot profile data generated by ImageJ.

#### Cell Preparation and Fluorescence-Activated Cell Sorting (FACS)

Retinas were dissected in AMES solution (Sigma A1420, equilibrated with 95% O2/5% CO_2_). Upon dissection, eyes and lenses were visually inspected for damage, blood, or inflammation, which were used as criteria for exclusion. Retinas were digested in papain and dissociated to single cell suspensions using manual trituration in ovomucoid solution. Cells were spun down at 450 g for eight minutes, resuspended in AMES+4% BSA to a concentration of 10 million cells per 100ml. 0.5ml of 0.2mg/ml anti-CD90.2-PE-Cy7 (Affymetrix eBioscience 25-0902-82) per 100ml of cells was incubated for 15 min, washed with an excess of media, spun down and resuspended again in AMES+4% BSA at a concentration of ∼7 million cells per 1 ml. Just prior to FACS the live cell marker Calcein Blue (Invitrogen C1429) was added. Cellular debris, doublets, and dead cells (Calcein Blue negative) were excluded. For FACS experiments with vGLUT2-Cre; LSL-Cas9 mice, RGCs were collected based CD90.2 and GFP double positive expression. For purification of transduced RGCs with CRISPR perturbation (sgRNAs co-expressed with mcherry), cells were collected based on CD90.2, GFP and mcherry triple positive expression. Cells were collected into ∼150ul of AMES+5% BSA.

#### Viscobead-induced experimental mouse glaucoma model

The elevation of intraocular pressure (IOP) was induced by injection of viscobeads to the anterior chamber of mouse eyes. The surgery procedures were modified by a well-established microbead occlusion model (Yang et al., 2012). Briefly, by using a standard double emulsion method, poly- d,l-lactic-co-glycolic acid (PLGA) / polystyrene (PS) core-shell microparticles (viscobeads) with 1- 20 um size distributed were first fabricated at a concentration of 30% (v/v) in saline. The corneas of anesthetized mice were gently punctured near the center using a 33g needle (CAD4113, sigma). A bubble was injected through this incision site into the anterior chamber to prevent the possible leakage. Then, 1 uL viscobeads were injected into the anterior chamber. After 5 min when the viscobeads were accumulated at the iridocorneal angle, the mouse was applied antibiotic Vetropolycin ointment (Dechra Veterinary Products, Overland Park, KS) and placed on a heating pad for recovery.

#### Intraocular pressure measurement

The IOP measurements were performed using a TonoLab tonometer (Colonial Medical Supply, Espoo, Finland) according to product instructions. Mice were first anesthetized with a sustained isoflurane flow (3% isoflurane in 100% oxygen). Average IOP was generated automatically with five measurements after the elimination of the highest and lowest values.

#### RNA-Seq library preparation

RNA from 5,000 – 10,000 FACS-sorted RGCs were isolated with RNeasy Micro Kit (Qiagen), and RNA-seq libraries were prepared with SMART-Seq v4 Ultra Low Input RNA Kit (Clontech), following manufacturers’ protocols. The cDNA was fragmented to 300 base pairs (bp) using the Covaris M220 (Covaris), and then the manufacturer’s instructions were followed for end repair, adaptor ligation, and library amplification. The libraries were quantified by the Qubit dsDNA HS Assay Kit (Molecular Probes); Library size distribution and molar concentration of cDNA molecules in each library were determined by the Agilent High Sensitivity DNA Assay on an Agilent 2200 TapeStation system. Libraries were multiplexed into single pool and sequenced using a HiSeq4000 instrument (Illumina, San Diego, CA) to generate 69 bp pair-end reads. The average uniquely-mapped, non-duplicated reads are ∼24 millions.

#### ATAC-Seq library preparation

ATAC-seq was performed as described with minor modifications (1). Briefly, 50,000 - 100,000 sorted RGCs were lysed in 100 µL ice-cold Resuspension Buffer (RSB, 10 mM Tris-HCl, 10 mM NaCl, 3 mM MgCl2, pH=7.4) with 0.1% NP40, 0.1% Tween- 20, and 0.01% Digitonin. Cell lysis were washed with 1 ml of cold ATAC-RSB containing 0.1% Tween-20 but NO NP40 or digitonin, and centrifuged at 4 °C for 10 minutes at 500 x g. Pelleted nuclei were resuspended gently in 50 µL transposition mix (25 ul 2x TD buffer, 2.5 ul transposase (100nM final), 16.5 ul PBS, 0.5 ul 1% digitonin, 0.5 ul 10% Tween-20, 5 ul H2O) and incubated for 30 minutes at 37 °C. DNA was cleaned using Zymo DNA Clean and Concentrator-5 Kit, and PCR amplified 5 cycles in a 50 µL reaction with Illumina Nextera adaptors using NEBNext High Fidelity 2x Master Mix. To determine the number of additional cycles to amplify the libraries, a side qPCR reaction was performed using 5 µL (10%) of the pre-amplified PCR. We used the Ct value of the qPCR at ¼ maximum fluorescence as the number of additional cycles. Final PCR products were cleaned using Zymo DNA Clean and Concentrator-5 Kit, and quantified by the KAPA Library Quantification kit prior to pooling and sequencing to an average depth of ∼90M unique non-mitochondria reads per sample on the Illumina NovaSeq platform at 2×100 bp.

#### Processing of RNA-seq data

RNA-seq data was processed and analyzed using custom pipeline (available at https://github.com/icnn/RNAseq-PIPELINE) as described previously (Norsworthy et al., 2017; Cheng et al., 2020). Raw sequenced reads were mapped to the reference genome Mus musculus (mm10) refSeq (refFlat) using STAR with default parameter (Dobin et al., 2013). Data quality was assessed on base-quality calls, nucleotide composition of sequences, insert sizes, per cent of uniquely aligned reads and transcript coverage using custom scripts and Picard Tools (http://broadinstitute.github.io/picard). More than 85% of reads were mapped uniquely to reference genome. Total counts of read fragments aligned to candidate gene regions were derived using the HTSeq program (https://htseq.readthedocs.io) with mouse mm10 refSeq (refFlat table) as a reference and used as a basis for the quantification of gene expression. Only uniquely mapped reads were used for subsequent analyses. Sequencing depth was normalized between samples using TMM in edgeR (Robinson et al., 2010). Outlier samples were removed using WGCNA (Langfelder and Horvath, 2008), with absolute sample connectivity score more than 2.5 standard deviation away from the mean. Genes with no counts in over 50% of all samples were removed.

#### RNA-seq analysis – differential gene expression

Principle component analysis (PCA) of the normalized expression data (first five PCs) was correlated with potential technical covariates, including experimental batch, aligning and sequencing bias calculated from STAR and Picard respectively. Differential expression analysis was conducted with the Bioconductor package edgeR (Robinson et al., 2010), including covariates that are significantly correlated with expression PCs: ∼ Genotype + AlignSeq.PC1 + AlignSeq.PC3 + AlignSeq.PC4. Statistical significance of differential expression was determined at FDR < 10% (q < 0.1).

#### Processing of ATAC-seq data

##### Alignment

Raw sequencing fastq files were assessed for quality, adapter content and duplication rates with FastQC, trimmed using trim-galore (https://github.com/FelixKrueger/TrimGalore) and aligned with bowtie2 (Langmead and Salzberg, 2012): bowtie2 --very-sensitive -X 2000 -x [reference_genome] -1 [input.left] -2 [input.right] | samtools view -hb -S - | samtools sort -o [sample_label].bam. The reference genome was mouse GRCm38 vM11. Samtools (Danecek et al., 2021) was used to calculate the read statistics, and Picard Tools (http://broadinstitute.github.io/picard) was used to remove duplicate.

##### ATAC-seq data QC - Transcription start site enrichment, fragment size distribution, chromHMM enrichment

Enrichment of open region signals at the transcription start site, an important quality control metric to evaluate ATAC-seq data, was measured by *tssenrich* (https://github.com/anthony-aylward/tssenrich). TSS positions were derived from RefSeq mm10, and the distance of each read within ± 2 kb centered on TSS were calculated. Read counts at each distance was summed up in a given library. A successful ATAC-seq library would form a characteristic shape with reads aggregated at the TSS, with an enrichment score > 7. Fragment size distribution is another quality control metric to visualize the nucleosome-sized periodicity resulting from chromatin digestion. This information was calculated by Picard Tools CollectInsertSizeMetrics. Enrichment of DA peaks within annotated genic regions of the genome or epigenetically annotated regions of the genome (Bogu et al., 2015) was calculated using the ratio between the (#bases in state AND overlap feature)/(#bases in genome) and the [(#bases overlap feature)/(#bases in genome) X (#bases in state)/(#bases in genome)] as we and others previously described (De La Torre Ubieta et al. 2018; Roadmap Epigenomics et al., 2015)

##### Peak calling and annotations

MACS2 (Zhang et al., 2008) was used to call peaks on each sample with the following command: macs2 callpeak -g mm --treatment [sample BAM] --format BAMPE --call-summits --nolambda --keep-dup all --min-length 100 -q 0.05. Peaks that overlapped with ENCODE mm10 blacklisted regions were removed. Peaks from each sample were merged to a set of union peaks across all conditions using bedtools merge (Quinlan et al., 2010). DiffBind (Stark et al., 2011) was used to obtain a consensus peak set by calculating the overlap rate. We kept peaks overlapped in at least two of the fifteen samples, at which point the overlap rate starts to drop off geometrically, indicating a good agreement among the peaksets. With this threshold, we obtained a total of 151,630 consensus peaks, with an average merged peak width of 1282 bp. The consensus peaks were further annotated to the transcriptional start site of genes in a distance of ± 2 kb from the gene start using ChIPseeker. Gencode vM24 was used for annotations.

##### ATAC-seq analysis - differential accessibility

We obtained the number of reads for consensus peaks (hereafter referred to as peaks) across samples using featureCounts from the Subread package (Liao et al., 2014), and the GC content using bedtools nuc. We then calculated normalization factors for each peak accounting for GC content, peak width, and total number of unique non-mitochondrial fragments sequenced using conditional quantile normalization from cqn package. We further performed principle component analysis (PCA) on cqn-normalized peak-count matrix using prcomp, and correlated technical and experimental variables to the first five PCs, including duplicate rate, fraction of reads in peaks (FRiP), TSS enrichment, batch, and injury conditions (Figure S2F). Batch effects significantly correlated with the peak-count PCs, which was then included as a covariate for differential accessibility analysis (Figure S2G). To obtain differential accessible peak-regions (DAR) comparing injured vs uninjured RGCs, we used a negative binomial regression with normalization based on the size factors from cqn (Hansen et al., 2012) and implemented in DEseq2 with default parameters (fitType=“parametric”, test=“Wald”) (Love et al., 2014). Normalized peak-count matrix with batch effects regressed out was used for subsequent analysis. MA-plot was used to visualize the general differential accessibility changes at 1- and 3-day following injury. Heatmaps and coverage plots of normalized reads within peaks were used to display DARs around specific genes, generated by ComplexHeatmap and Gviz respectively.

##### ATAC-seq analysis – linking proximal and distal regulatory elements to cognate genes

Using GENCODE annotations, we defined an ATAC-seq peak ± 2 kb of a gene’s transcription start site (TSS) as a promoter (proximal regulatory element), and non-promoter peaks ± 500 kb of TSS as distal regulatory regions for that gene. To correlate differential accessible promoter/distal peaks to gene expressions, we computed the sample-wise Pearson’s correlations between normalized read counts from ATAC-seq and RNA-seq data. For genes with multiple distal regulatory links in the ± 500 kb window, the average accessibility of distal regulatory elements was used to correlate with gene expression. Next, to account for spurious associations, we generated a background null model by computing correlations between randomly selected peaks and randomly selected genes on different chromosomes. We calculated the mean and standard deviation of this null distribution of correlations, enabling us to compute p-values for the test correlations. The correlative peak-gene pairs with FDR < 0.1 were further clustered by *K*- means and genes linked each cluster were annotated with gprofiler2 for general biological pathways in Gene Ontology (GO). GO terms were chosen based on their FDR-corrected p-values and relevance to the current study.

##### ATAC-seq analysis – transcription factor motif activity

To find TF motifs within peaks, we used motif position weight matrices (PWMs) integrated from JASPAR2016, HOCOMOCCO v10, UniPROBE and SwissRegulon, which include a core set of 1,312 non-redundant motifs from human and mouse (Funk et al., 2020). This integrated PWMs were used to scan differential accessible regions for motif occurrence by motifmatchr, resulting in a binary peak-by-matches matrix. For each motif, we computed the odds ratio and the significance (P-value < 5×10^−5^) of enrichment comparing to background nucleotide frequencies across input peak regions using Fisher’s exact test. The degree of accessibility at enriched TF motifs across samples was computed by chromVAR (Schep et al., 2017), which quantitatively measures changes of ATAC-seq counts in peaks containing the TF motifs as deviation Z-scores. To correspond TF motifs to TF genes, ChromVAR deviation z-scores for each TF motif were correlated to the TF gene’s log2- transformed TPM values across samples. If the correlation between motif and gene expression is greater than 0.5 with an adjusted p-value less than 0.05 and a maximum cross-sample difference in deviation z-score that is in the top quartile, the TF is classified as a positive regulator of chromatin state; if the correlation is less than – 0.5 with an adjusted p-value less than 0.05 and a maximum deviation score in the top quartile, it is classified as a negative TF regulator of chromatin state. To visualize, we plotted the TF gene – motif correlations against the maximum cross-sample difference in deviation z-score for all TFs, with the top ‘TF hits’ meeting the criteria of positive or negative TF regulators of chromatin state described above.

##### ATAC-seq analysis – transcription factor footprinting

###### Tn5 bias correction and footprinting

The first step of fooptrinting is to correct Tn5 transposase cleavage bias in the ATAC-seq data. To do this, we first merged biological replicates from each condition by Picard Tools and downsampled ATAC-seq BAMs to a depth of 60 million reads using samtools. TOBIAS (Bentsen et al., 2020) ATACorrect module was applied to merged, down- sampled reads within the consensus peaks to estimate the background bias of Tn5 transposase. Subtracting the background Tn5 insertion cuts from the uncorrected signals yields a corrected track, highlighting the effect of protein binding. The footprint score was calculated by TOBIAS ScoreBigWig, which measures both accessibility and depth of the local footprint, thus correlating with the presence of a TF at its target loci, and the chromatin accessibility of the regions where this TF binds.

###### Assigning TFs to footprints

To match footprints to potential TF binding sites, and to estimate TF binding activity on its target loci, we applied TOBIAS BINDetect module to the corrected ATAC-seq signals within peaks, with the same set of TF motif PWMs used for ChromVAR as input. This method obtains the positions of TF binding sites, which are then mapped to the footprints for each condition. Each footprint site was assigned a log2FC (fold change) between two conditions, representing whether the binding site has larger/smaller TF footprint scores in comparison. To calculate statistics, a background distribution of footprint scores is built by randomly subsetting peak regions at ∼200bp intervals, and these scores were used to calculate a distribution of background log2FCs for each comparison of two conditions. The global distribution of log2FC’s per TF was compared to the background distributions to calculate a differential TF binding score, which represents differential TF activity between two conditions. A P-value is calculated by subsampling 100 log2FCs from the background and calculating the significance of the observed change. By comparing the observed log2FC distribution to the background log2FC, the effects of any global differences due to sequencing depth, noise etc. are controlled. To visualize, we used soGGI to plot TF footprints, and bar graphs to show the global TF footprint activity changes comparing injured to uninjured RGCs.

###### Linking TF footprint sites to its targeted genes combining RNA-seq data

Footprint sites which are bound by each of the four TF with increased footprint score after injury were considered as injury-responsive TF-footprints. Distribution of these footprints were further annotated with ChIPseeker (Yu et al., 2015), and the footprints located within ± 500 kb of a gene’s TSS were linked to that gene. We did not consider distal intergenic regions that are > ± 500 kb of TSS as linking them to a gene requires higher-order chromatin conformation data. Genes that are footprinted by the TF and differentially regulated in the RNA-seq upon CRISPR ablation of this TF are considered as direct target genes of each TF. The similarity and dis-similarity of each TF’s target genes were analyzed by calculating sample distances and then hierarchically clustered. Genes that are unique or common to each TF were annotated with gprofiler2 and Ingenuity Pathway Analysis (IPA) Software (Qiagen) for gene ontology analysis. GO terms were chosen based on their FDR-corrected p-values and relevance to the current study.

### QUANTIFICATION AND STATISTICAL ANALYSIS

The normality and variance similarity were measured by Microsoft Excel and R programming Language before we applied any parametric tests. Two-tailed student’s t-test was used for the single comparison between two groups. The rest of the data were analyzed using one-way or two-way ANOVA depending on the appropriate design. *Post hoc* comparisons were carried out only when the primary measure showed statistical significance. P-value of multiple comparisons was adjusted by using Bonferroni’s correction. Error bars in all figures represent mean ± S.E.M. The mice with different litters, body weights and sexes were randomized and assigned to different treatment groups, and no other specific randomization was used for the animal studies.

## SUPPLEMENTAL INFORMATION

Supplementary information includes 7 supplementary figures and 6 supplemental tables.

## AUTHOR CONTRIBUTIONS

We thank A. Nazzari, R. Wang and A. Zhu for assistance of library preparation. F.T., Y.C., J.R.S., D.H.G. and Z.H. conceived and F.T., Y.C., S.Z., Q.W., A. M., K.G., W. J., R.K., Q.W., Q.W., M.T., R.D., H.M., A.J., S.H., J.Y., X.C., S.H., J.S., P.D., D.T. performed the experiments and/or analyzed the data. F.T. performed sample collection. Y.C. and K.G. performed ATAC-seq library preparation. Q.W. and F.T. performed RNA-seq library preparation. Y.C. implemented analysis pipeline for ATAC-seq data and performed integration with RNA-seq data. R.K. pre-processed RNA-seq data. Y.C., F.T., and W.J. performed downstream analysis. F.T., Y.C., J.R.S., D.H.G. and Z.H. prepared the manuscript with the inputs from all authors.

## ACKNOWLEDGEMENTS

This study was supported by grants K99 EY032181 to F.T., EY030204-01 to J.R.S and Z.H., R01EY021526 and R01EY026939 to Z.H., Wings for Life Spinal Cord Research Foundation to Y.C. and A.J., and the Dr. Miriam and Sheldon G. Adelson Medical Research Foundation (to C.J. W., D.H.G. and Z.H., Gilbert Family Foundation (to Z.H.). IDDRC and viral cores were supported by the grants from the NIH (HD018655 and P30EY012196).

## SUPPLEMENTAL INFORMATION

**Table S1.** ATAC-seq metadata, quality control metrics, and annotation of differentially accessible regions (DARs) comparing RGCs injured at day 1 or 3 versus uninjured condition (day 0). Related to Figure 2. (A) ATAC-seq metadata and quality control metrics including percent of duplicated reads after sequencing (DuplicateRate), non-duplicated, uniquely mapped reads (UniqMappedReads), total number of peaks from MACS2 (TotNarrowPeak), TSS enrichment, and fraction of reads in peaks (FRiP). Annotation of injured RGC DARs [absolute log fold changes (logFC) > 0.3 and false discover rate (FDR) < 0.1] comparing day 1 *v.s.* day 0 (B) or day 3 *v.s.* day 0 (C).

**Table S2.** RNA-seq metadata, quality control metrics, and annotation of differentially expressed genes (DEGs) comparing injured RGCs at indicated time points versus uninjured ones (day 0). Related to Figure 2. (A) RNA-seq metadata and quality control metrics obtained from *Picard Tools*. Principle component analysis (PCA) was performed on the metrics and the first seven PCs accounting > 95% of variance was kept for further analysis. (B-C) Annotation of injured RGC DEGs (absolute logFC > 0.3 and FDR < 0.1) comparing day 1 *v.s.* day 0 (B) or day 3 *v.s.* day 0 (C).

**Table S3.** Clustering and annotation of putative peak-gene links. Related to Figure 2. (A) Linked peak-gene pairs. Proximal or distal peaks that are differential accessible are linked to corresponding genes that are differentially expressed at day 1 or day 3 following optic nerve crush. The peak-gene pairs are grouped into two clusters: the ‘up’ cluster where the accessibility of linked peaks and the expression of corresponding genes are both up-regulated, and the ‘down’ cluster where both are down-regulated. (B) Gene Ontology of the up-regulated cluster. (C) Gene Ontology of the down-regulated cluster.

**Table S4.** TF motif activity in injured RGCs, related to Figure 3. (A) ChromVAR-estimated motif matches in DARs. A list of non-redundant, human or mouse TF motifs (Methods) was used, named in the format of [species]-[database]-[corresponding TF]-[motif ID]. ChromVAR returns variability, which is the standard deviation of the motif z-scores computed across samples, and boostrap confidence intervals and *p*-value for the variability by resampling samples. These calculations are correlative to motif accessibility changes across samples. (B) Sample-wise correlations of motif activity to the expression levels of corresponding TF.

**Table S5.** RNA-seq metadata, quality control metrics and annotation of differentially expressed genes comparing RGCs with or without CRISPR ablation of one of the 4 survival TFs (ATF3, ATF4, C/EBPγ CHOP). Related to Figure 4-5. (A). RNA-seq metadata and quality control metrics obtained from *Picard Tools*. (B-E) Annotation of DEGs in RGCs with ablation of ATF3, ATF4, C/EBPγ, CHOP, respectively.

**Table S6.** Target genes regulated by ATF3/CHOP, or ATF4/C/EBPγ or by all 4 TFs predicted by DNA-footprinting and RNA-seq. Related to Figure 4-5. (A) Genes that are positively or negatively regulated by ATF3/CHOP, but not ATF4/C/EBPγ. (B) Genes that are positively or negatively regulated by ATF4/C/EBPγ, but not ATF3/CHOP. (C) Genes that are positively or negatively regulated by all four TFs.

---

**Figure S1.**
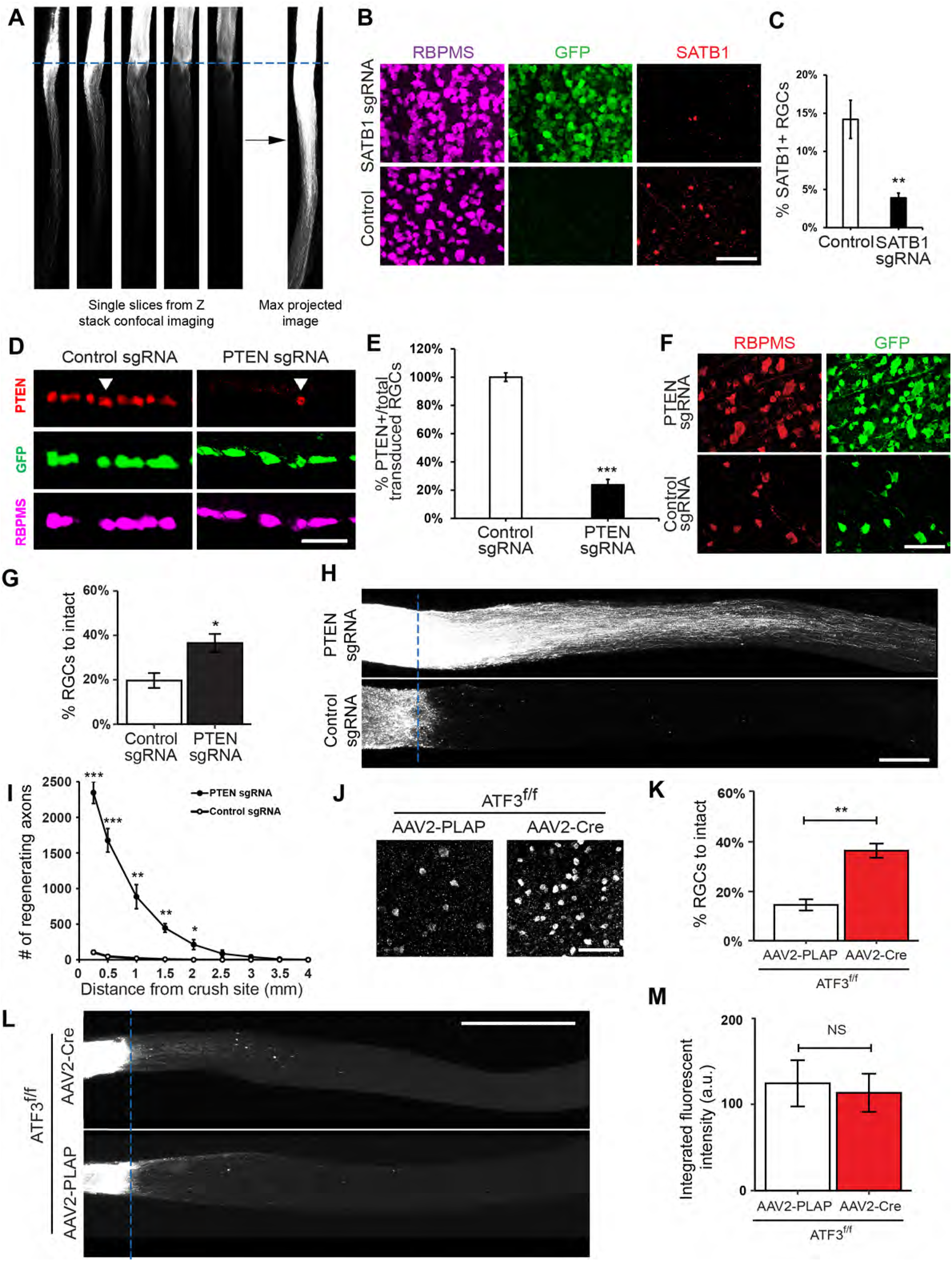
Characterization of in vivo CRISPR knockout in ONC model, Related to Figure 1. (A) Identification of crush site from wholemount optic nerves after iDISCO tissue clearing. (B-C) Knockout efficiency of SATB1 sgRNA injection, including immunohistochemistry images (B) and quantification (C). (D-E) Knockout efficiency of PTEN sgRNA injection, including immunohistochemistry images (D) and quantification (E). Data are shown as mean ± s.e.m. with n = 4-6 biological repeats. Scale bar: 50 μm. (F-G) RGC survival phenotype of PTEN sgRNA injection after ONC injury, including immunohistochemistry images (F) and quantification (G). Data are shown as mean ± s.e.m. with n = 4-6 biological repeats. Scale bar: 50 μm. (H-I) Axon regeneration phenotype of PTEN sgRNA injection after ONC injury, including CTB fluorescent imaging (H) and quantification (I). Data are shown as mean ± s.e.m. with n = 4-6 biological repeats. Scale bar: 0.5 mm. (J-M) RGC survival and axon regeneration phenotypes in ATF3 *f/f* mice intravitreally injected with AAV2-Cre and ONC. (J) Representative immunohistochemistry images of retinal sections showing the RGC survival phenotype and its quantification (K). Data are shown as mean ± s.e.m. with n = 4-6 biological repeats. Scale bar: 50 μm. (L) Representative CTB fluorescent images of optic nerves showing the axon regeneration phenotype and its quantification (M). Data are shown as mean ± s.e.m. with n = 4-6 biological repeats. Scale bar: 0.5 mm.

**Figure S2.**
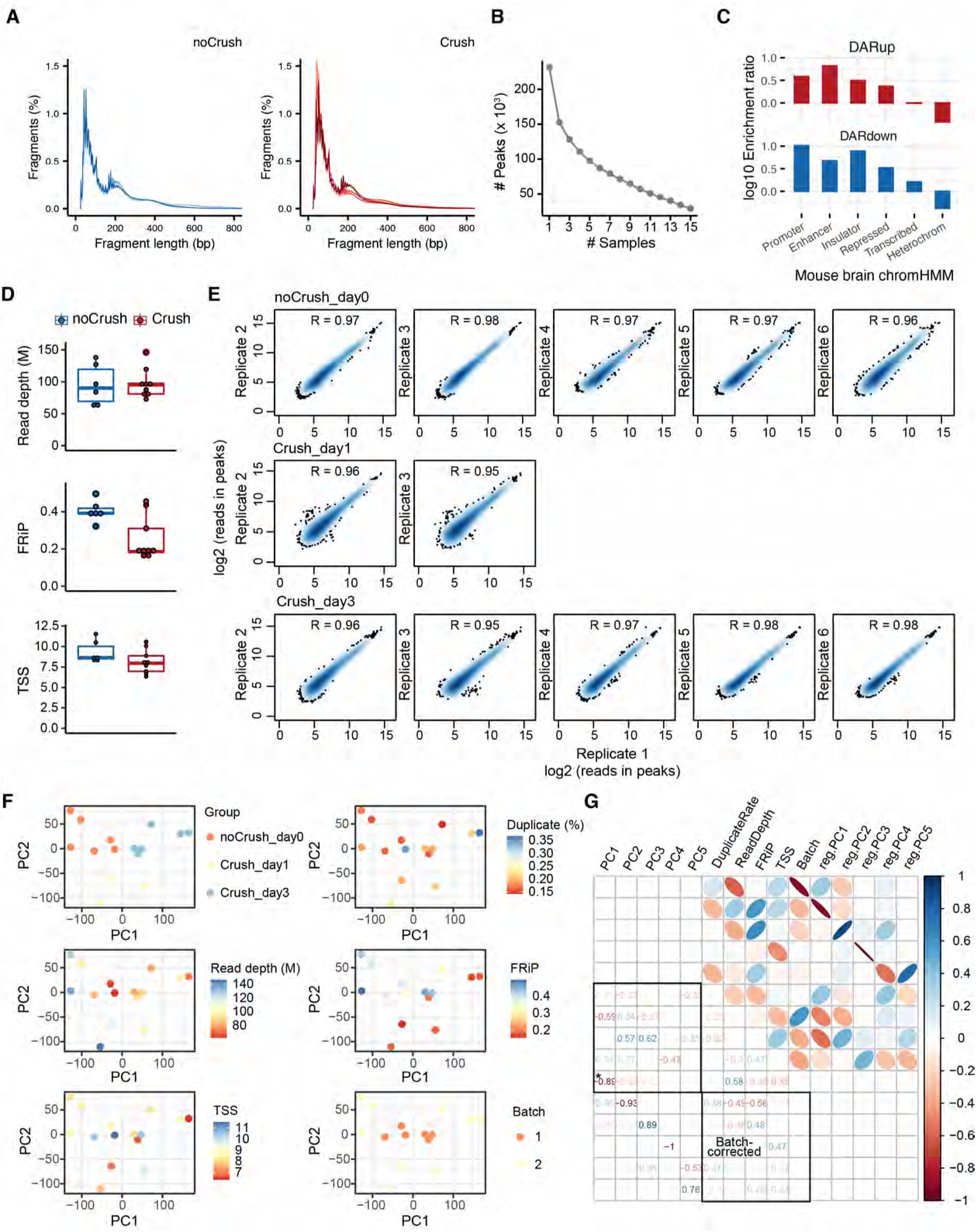
ATAC-seq data quality measures, Related to Figure 2. (A) Insert size histogram showing the expected pattern of transposase insertion for individual samples. (B) Number of overlapping peaks (y-axis) in the number of samples indicated (x-axis). A geometric drop-off of overlapped peaks was expected when the number of samples increases. A minimum overlap of 2 samples was used to derive consensus peak sets as the drop-off rate begins to stabilize when N=2. (C) Enrichment of differentially accessible regions with more within annotated genomic regions from mouse brain. DARup: more accessible after injury at day 1 or 3. DARdown: less accessible after injury at day 1 or 3. (D) Other important ATAC-seq data quality metrics including the number of uniquely mapped non-mitochondria reads, fraction of reads in peaks (FRiP) and TSS enrichment for individual samples. (E) Biological RGC replicates of same conditions also show high correlation of normalized reads in consensus peaks. (F) Principle component analysis of sequence-dept, GC-content normalized counts in consensus peaks. Colors indicate the biological and technical covariates of each sample, including group, duplicate rate, uniquely mapped reads, fraction of reads in peaks, TSS enrichment, and batch. (G) Pearson’s correlation heatmap between experimental and technical variables and the first five PCs of peak-count matrix prior (PC1-5) or after batch correction (reg.PC1-5).

**Figure S3.**
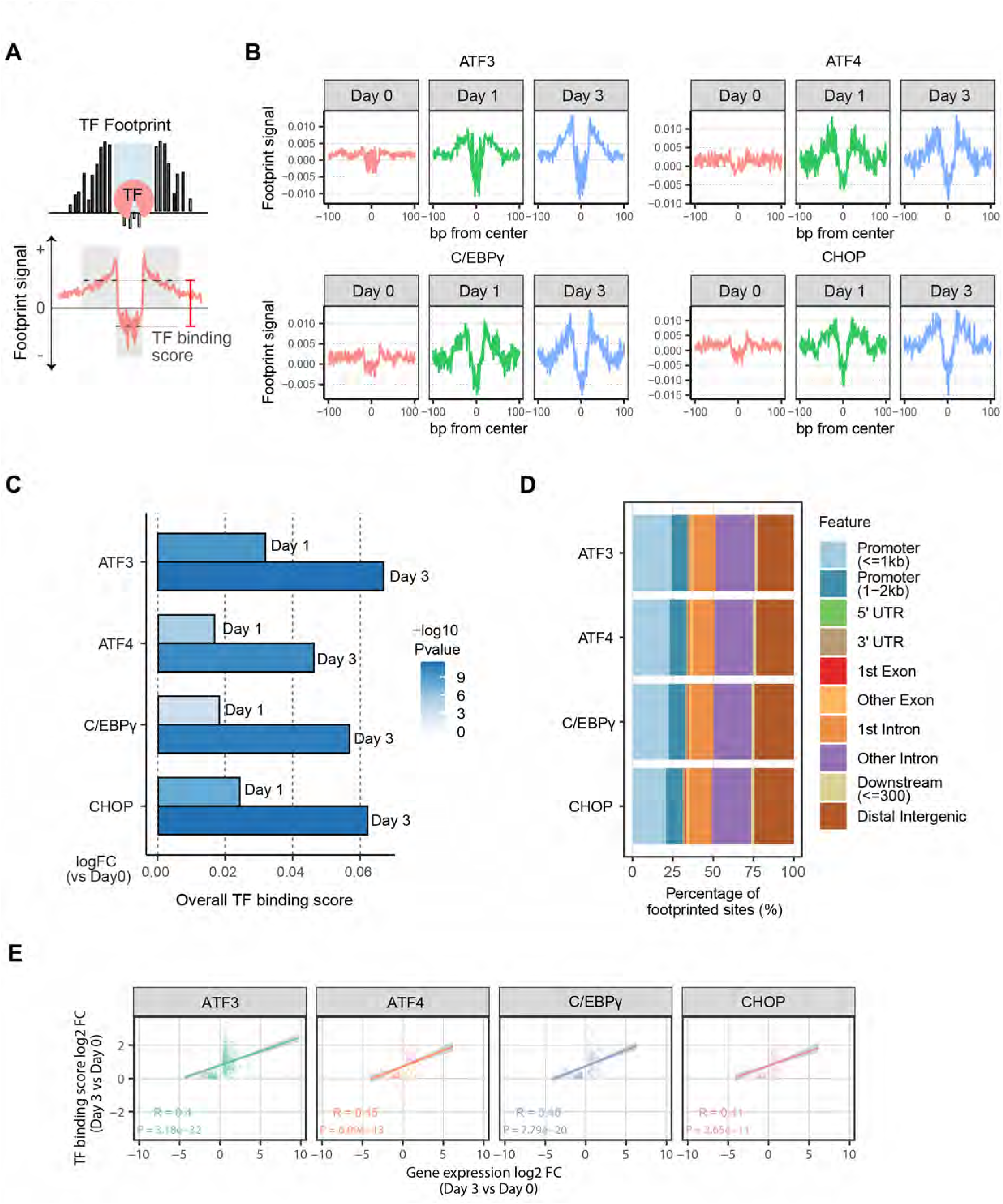
DNA footprinting to identify direct binding sites of the four survival TFs in injured RGCs, Related to Figure 3. (A) A schematic diagram demonstrating typical footprint patterns of regions bound by individual TFs. Following Tn5 insertion bias correction and TF-footprint detection, a TF binding score measuring both accessibility and depth of each footprint was calculated, correlating with the presence of a TF at its target loci, and the chromatin accessibility of the regions where this TF binds. (B) Tn5-bias-corrected footprint signals centered around each TF’s binding sites. (C) Differential TF binding score for each TF. Each footprint site was assigned a log2FC (fold change) per comparison (day1 vs day 0, or day 3 vs day 0), representing whether the binding site has larger/smaller TF binding scores in comparison. The global distribution of log2FCs per TF is compared to the background distributions to calculate a differential TF activity score. A p-value is calculated by subsampling 100 log2FCs from the background and calculating the significance of the observed change, shown as the bar colors. (D) Genome distribution of each TF’s footprints. (E) Pearson correlations between differential TF binding scores of footprints and differential gene expression levels of linked genes.

**Figure S4.**
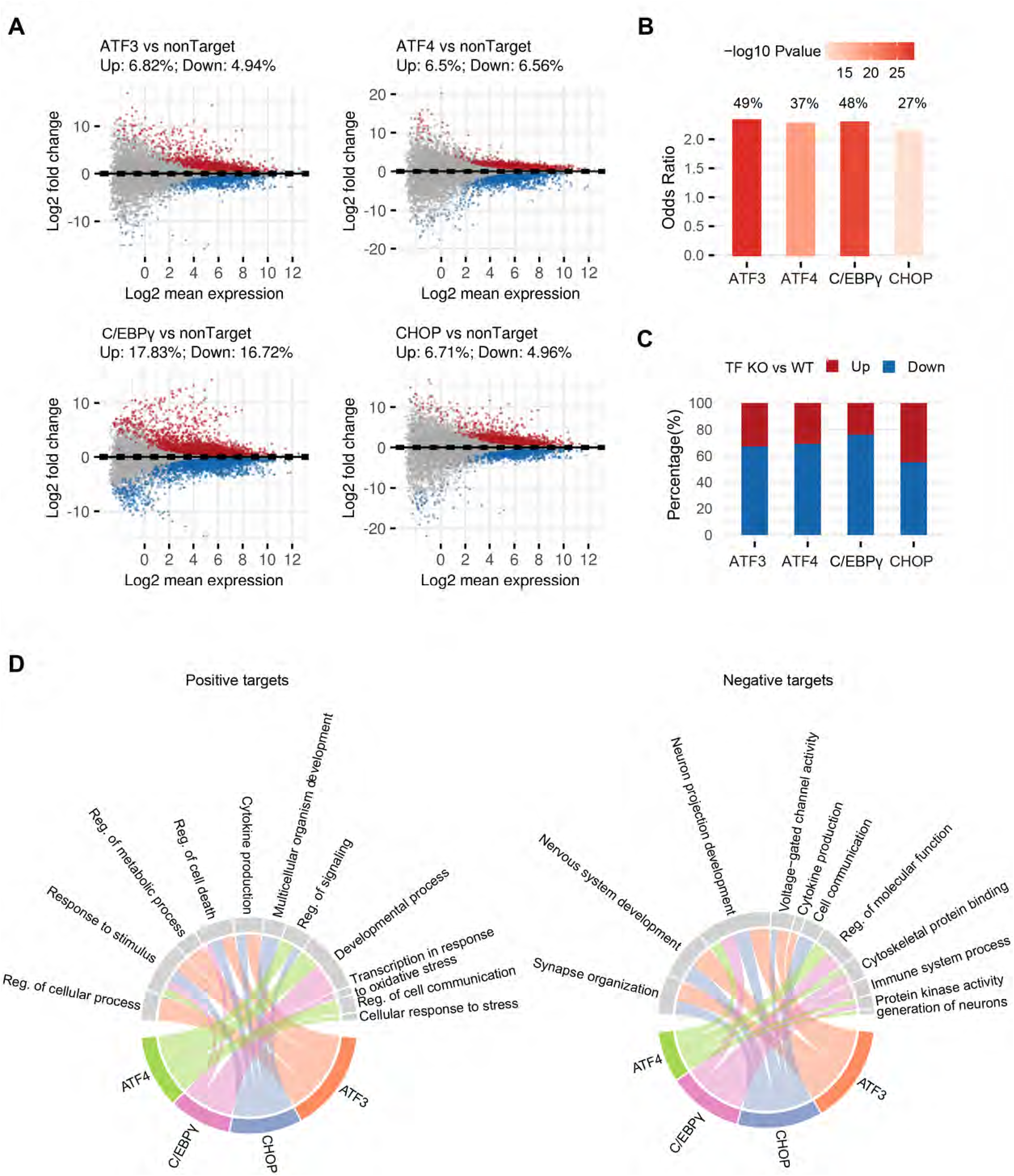
Interrogation of the results from TF-footprinting and RNA sequencing of RGCs with or without TF knockout, Related to Figure 4. (A) MA plots from means to log fold changes displaying differential expressed genes (DEGs) in the RNA-seq of injured RGCs (at day 3) bearing CRISPR ablation of individual TF. Each dot represents a gene, and colored dots indicate DEGs (FDR < 0.1, | log2 FC| > 0.3). Up-regulated: red; Down-regulated: blue. (B) Overlap between genes predicted to be targeted by each TF by footprint analysis and the genes differentially regulated in the RNA-seq upon TF CRISPR ablation. Fisher’s exact test was used to test the odds ratio and p-value of the overlap. Numbers shown above each odds ratio bar represents the percent of footprinted genes overlapping with DEGs in the RNA-seq. (C) Percentage of up- or down-regulated genes directly targeted by each TF. (D) Chord diagram showing the top general biological pathways associated with each TF’s positive or negative targets. Link width indicates log10 (enrichment FDR p-values) of the GO term. For positive targets, the FDR p-values range from 1.19e-02 ∼ 5.11e-12; for negative targets, the FDR p-values range from 1.49e-03 ∼ 7.87e-18.

**Figure S5.**
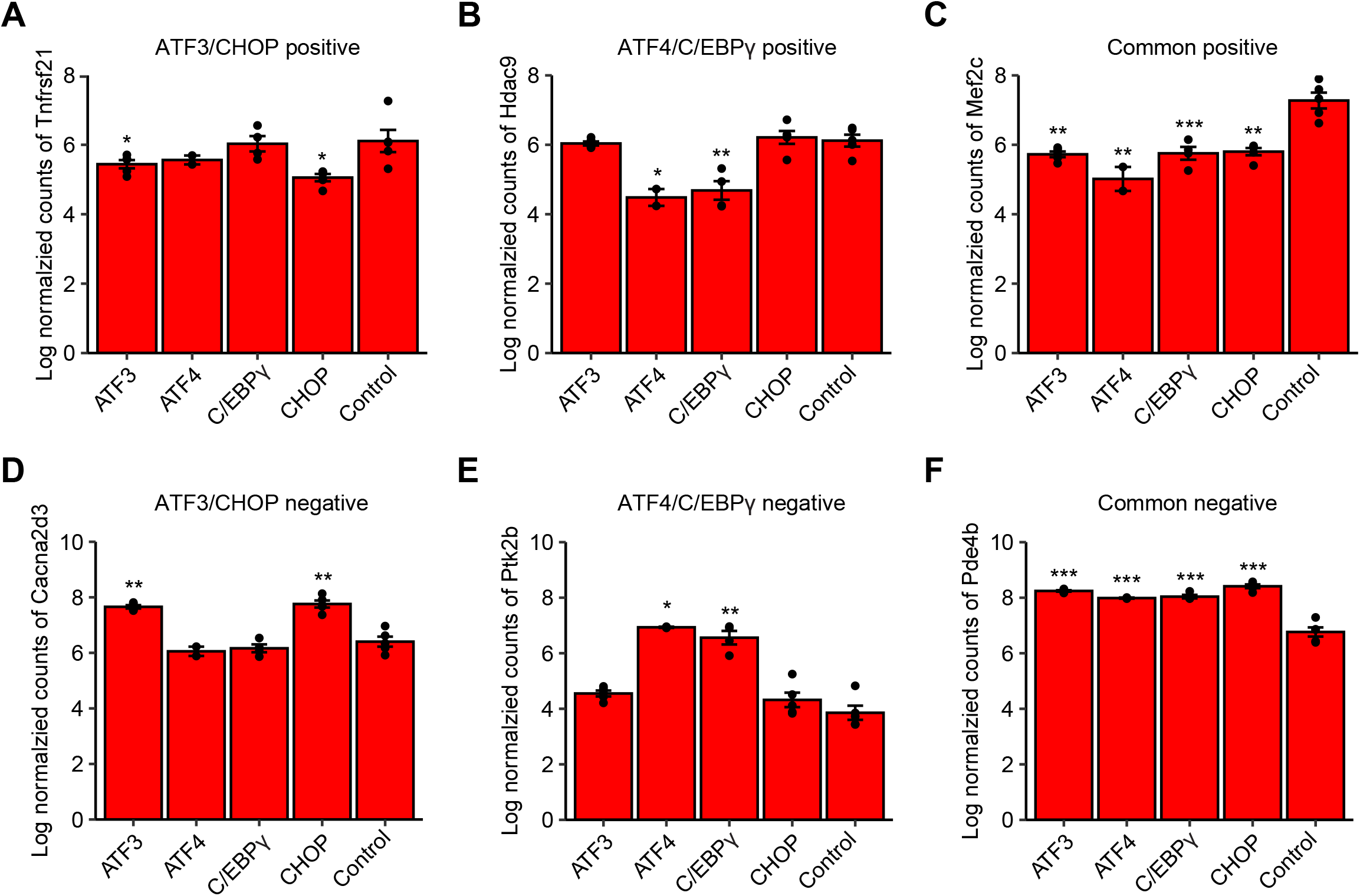
Verification of the two injury-induced transcriptional programs by perturbed RNA-seq, Related to Figure 5. (A-F) Normalized RNA-seq counts in log-scale for representative genes shown in Figure 5 under each perturbed RNA-seq conditions (nontargeting, ATF3 KO, ATF4 KO, C/EBPγ KO and CHOP KO). *FDR < 0.1; **FDR < 0.05; ***FDR < 0.01.

**Figure S6.**
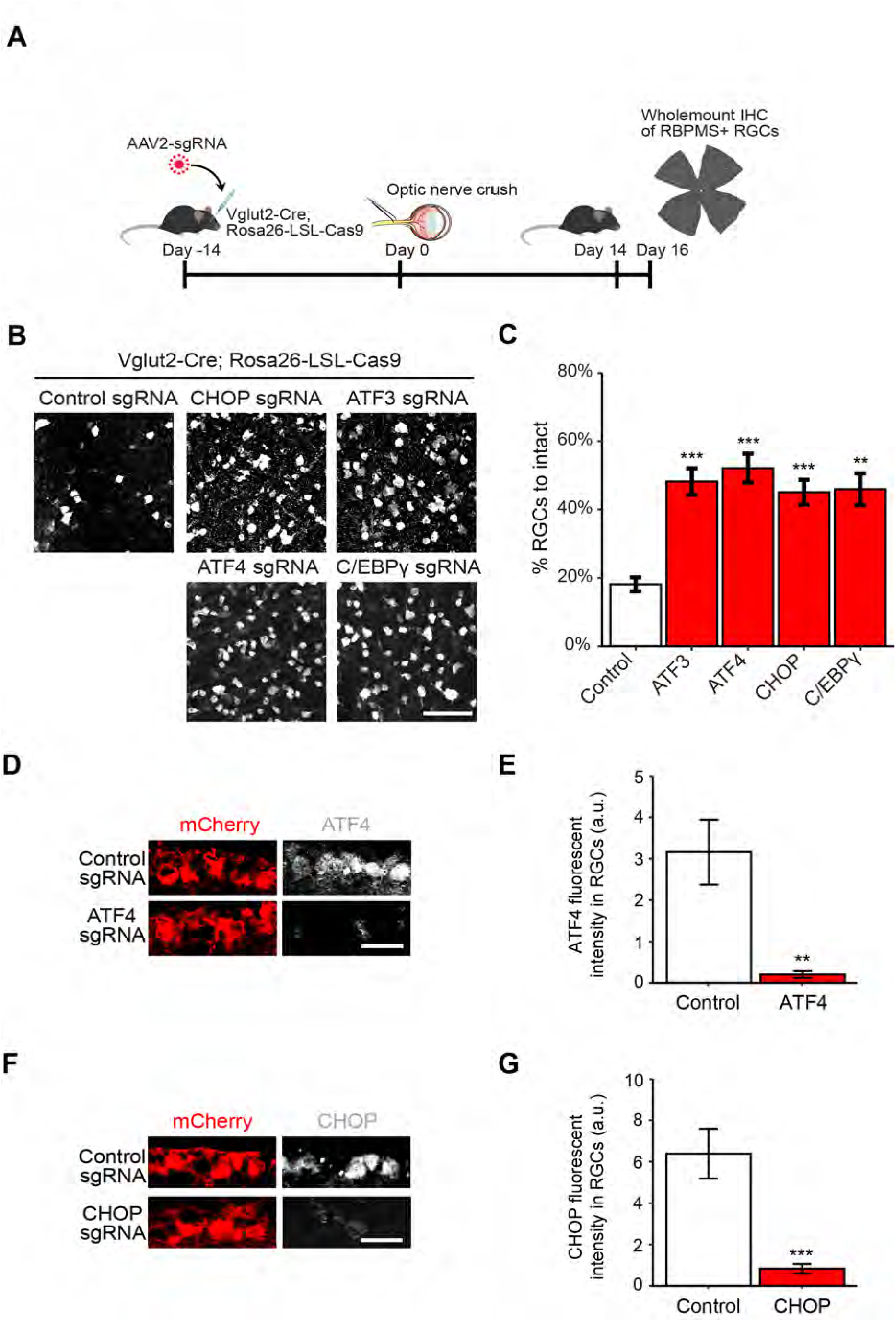
Functional characterization of the key TFs, Related to Figure 6. (A-C) Verifying cell autonomous effects of ATF3/ATF4/C/EBPγ/CHOP. (A) Schematic of RGC specific ablation for genes based in vGlut2-Cre; LSL-Cas9 mice. (B) Representative images of retinal sections showing RGC survival with RGC-specific CRISPR ablation. (C) Quantification of the RGC survival with RGC-specific CRISPR knockout based on (B). Data are shown as mean ± s.e.m. with n = 4-6 biological repeats. *p<0.05, **p<0.01, ***p<0.001. Scale bar, 50 μm. (D-E) Representative retinal images (D) and quantification (E) of ATF4 immunostaining indicate the knockout efficiency of sgRNAs targeting the ATF4. Data are shown as mean ± s.e.m. with n = 4. **p<0.01. Scale bar, 10 μm. (F-G) Representative retinal images (F) and quantification (G) of CHOP immunostaining indicate the knockout efficiency of sgRNAs targeting the CHOP. Data are shown as mean ± s.e.m. with n = 4. ***p<0.001. Scale bar, 10 μm.

**Figure S7.**
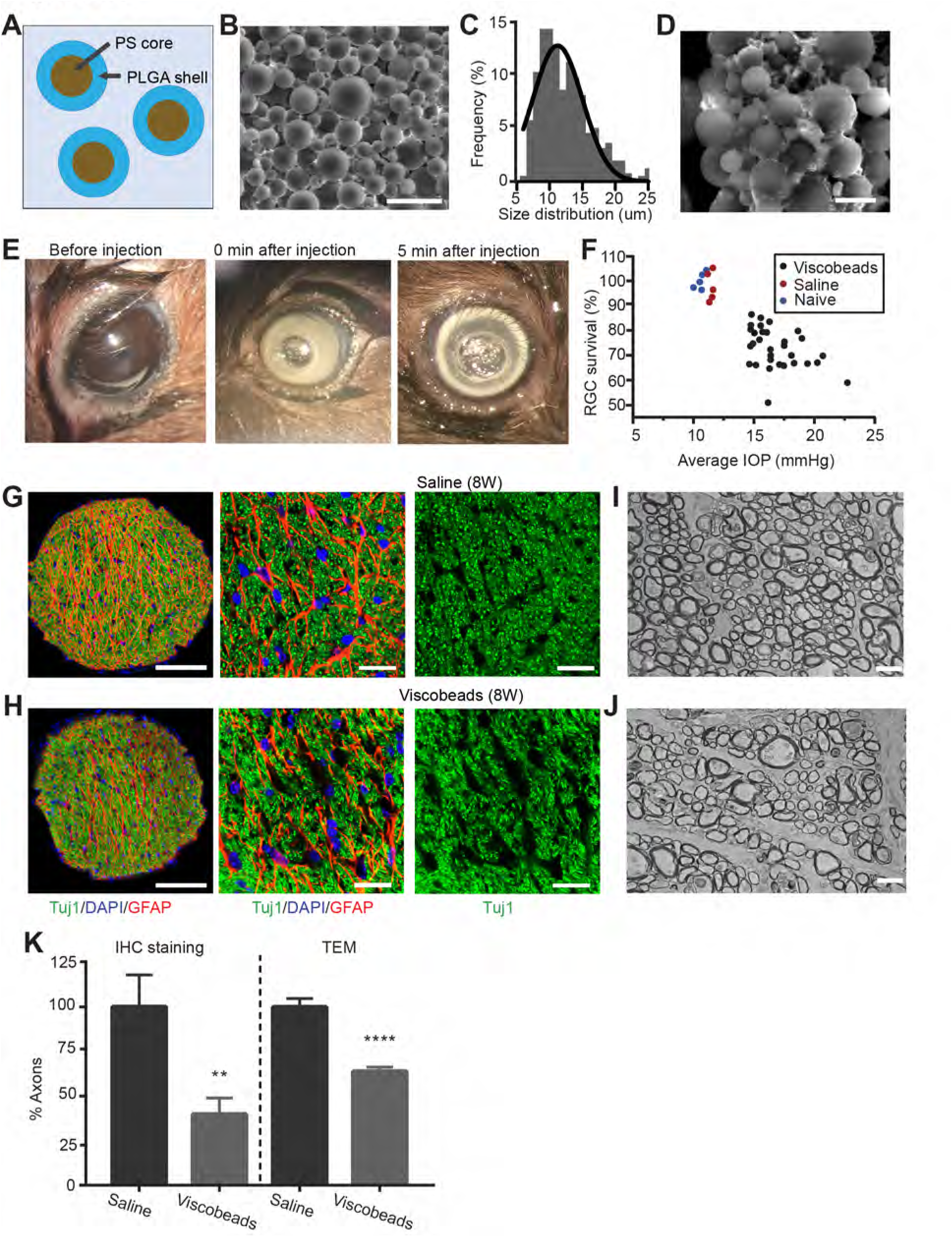
The viscobeads occlusion experimental glaucoma model, Related to Figure 7. (A) Schematic illustration of viscobeads with core-shell structures, with biodegradable PLGA shell as viscoelastic materials with gradual degradation after injection and non-degradable PS core as the long-term mechanical clogging. (B) A representative scanning electron microscope (SEM) image of viscobeads. Scale bar, 50 μm. (C) The size distribution of viscobeads. (D) The SEM images of viscobeads before (left) and 8 weeks after injection (right). The degraded PLGA became viscoelastic materials and glued the individual beads. Scale bar, 10 μm. (E) Representative photographs of a mouse eye before and after injection. The white viscobeads were injected into the anterior chamber (middle), then became accumulated at iridocorneal angle 5 min after injection (right). (F) The distribution of average IOP with RGC survival for 8 weeks after the viscobeads (n = 31), or saline (n = 5) injection, and the naïve group (n = 5). Data are shown as mean ± SD. (G-K) Representative confocal images (G, H), TEM images (I, J), and quantification of axon numbers (K) of optic nerve cross-sections from the mice with saline (G, I) or viscobeads (H, J). Scale bar for the left panel of (G, H), 100 μm; middle and right panels of (G, H), 20 μm. Scale bar for (I, J), 2 μm.

## Notes

### Competing Interest Statement

C.J.W. is a founder of Nocion Therapeutics
and QurAlis.
J.R.S. is a consultant for Biogen.
Z.H. is an advisor of SpineX, Life Biosciences and Myro Therapeutics.

